# Structures of the human mitochondrial ribosome recycling complexes reveal distinct mechanisms of recycling and antibiotic resistance

**DOI:** 10.1101/2020.12.20.423689

**Authors:** Ravi Kiran Koripella, Ayush Deep, Ekansh K. Agrawal, Pooja Keshavan, Nilesh K. Banavali, Rajendra K. Agrawal

## Abstract

Ribosomes are recycled for a new round of translation initiation by dissociation of ribosomal subunits, messenger RNA and transfer RNA from their translational post-termination complex. Mitochondrial ribosome recycling factor (RRF_mt_) and a recycling-specific homolog of elongation factor G (EF-G2_mt_) are two proteins with mitochondria-specific additional sequences that catalyze the recycling step in human mitochondria. We have determined high-resolution cryo-EM structures of the human 55S mitochondrial ribosome (mitoribosome) in complex with RRF_mt_, and the mitoribosomal large 39S subunit in complex with both RRF_mt_ and EF-G2_mt_. In addition, we have captured the structure of a short-lived intermediate state of the 55S•RRF_mt_•EF-G2_mt_ complex. These structures clarify the role of a mitochondria-specific segment of RRF_mt_ in mitoribosome recycling, identify the structural distinctions between the two isoforms of EF-G_mt_ that confer their functional specificity, capture recycling-specific conformational changes in the L7/L12 stalk-base region, and suggest a distinct mechanistic sequence of events in mitoribosome recycling. Furthermore, biochemical and structural assessments of the sensitivity of EF-G2_mt_ to the antibiotic fusidic acid reveals that the molecular mechanism of antibiotic resistance for EF-G2_mt_ is markedly different from that exhibited by mitochondrial elongation factor EF-G1_mt_, suggesting that these two homologous mitochondrial proteins have evolved diversely to negate the effect of a bacterial antibiotics.

## Introduction

The process of protein synthesis in all living cells is orchestrated by highly complex macromolecular assemblies called ribosomes, in coordination with mRNA, tRNAs and multiple translational factors. Mitochondrial ribosomes (mitoribosomes) and their associated translation machinery are distinct from those in the cytoplasm and display features reminiscent of prokaryotic translation (Pel and Grivell, 1994), in line with the assumption that mitochondria have evolved from endocytosis of an α-proteobacterium by an ancestral eukaryotic cell (Gray et al., 1999). However, cryo-electron microscopy (cryo-EM) structures have revealed that the mammalian mitoribosomes have diverged considerably from their bacterial counterparts and acquired several unique features (Amunts et al., 2015; Brown et al., 2014; Desai et al., 2017; Greber et al., 2015; Kaushal et al., 2014; Koripella et al., 2020; Koripella et al., 2019b; Sharma et al., 2003). A striking difference is the reversal in the protein to RNA ratio, as the bacterial ribosomes are high in ribosomal RNA (rRNA) whereas the mammalian mitoribosomes are high in protein. The increase in protein mass is the result of acquisition of multiple mito-specific ribosomal proteins (MRPs) and addition of extensions to many MRPs that are homologous to bacterial ribosomal proteins. Though the steps of mitochondrial translation closely resemble those for prokaryotic translation in the general sequence of events and the homologous accessory protein factors involved, they also show significant structural and functional differences (Christian and Spremulli, 2012; Sharma et al., 2013).

The complex process of protein synthesis is accomplished in four essential steps of initiation, elongation, termination and the ribosome recycling. Transitioning from translation termination to ribosome recycling has been best characterized in eubacteria. During translation termination, the nascent polypeptide chain attached to the peptidyl tRNA is released from the ribosome with the help of a class I release factor (RF) that interacts with the stop codon exposed at the ribosomal decoding site, or aminoacyl-tRNA binding site (A site) (Ito et al., 1996; Kisselev et al., 2003; Poole and Tate, 2000). Subsequently, the class I RF is dissociated from the ribosome with the help of a class II RF in a GTP hydrolysis-dependent manner (Freistroffer et al., 1997; Gao et al., 2007; Zavialov et al., 2002). At the end of the termination, the translated mRNA and the deacylated tRNA remain associated with the ribosome (Kaji et al., 2001; Karimi et al., 1999), a state referred to as the post-termination complex (PoTC). In order to initiate a new round of protein synthesis, the ribosome must be split into its two subunits and its bound ligands must be removed. In eubacteria, the disassembly of the PoTC requires the concerted action of two protein factors, the ribosome recycling factor (RRF) and the elongation factor G (EF-G) (Ito et al., 2002; Janosi et al., 1998; Kaji et al., 2001; Rao and Varshney, 2001; Zavialov et al., 2005). RRF binds to the PoTC as the 70S ribosome adopts a ratcheted conformation (Barat et al., 2007; Gao et al., 2005), in which the small (30S) subunit of the ribosome rotates in an anticlockwise direction with respect to the large (50S) subunit (Frank and Agrawal, 2000). This is followed by the binding of EF-G in conjugation with guanosine 5’-triphosphate (GTP) to the RRF-bound PoTC and the dissociation of the 70S ribosome into its large and small subunits, a process that requires the hydrolysis of GTP on EF-G (Borg et al., 2016; Hirokawa et al., 2005; Peske et al., 2005; Zavialov et al., 2005). Though the involvement of a third factor, initiation factor 3 (IF3) in the recycling process is generally agreed upon, its precise function has been debated (Hirokawa et al., 2005; Iwakura et al., 2017; Karimi et al., 1999; Zavialov et al., 2005).

Unlike eubacteria, where a single form of EF-G participates in both the elongation and ribosome recycling steps (Borg et al., 2016; Savelsbergh et al., 2009), mammalian mitochondria utilize two isoforms of EF-G, EF-G1_mt_ and EF-G2_mt_ (Hammarsund et al., 2001; Tsuboi et al., 2009). While EF-G1_mt_ specifically functions as a translocase during the polypeptide elongation step (Bhargava et al., 2004), EF-G2_mt_ has been reported to act exclusively as a second recycling factor together with RRF_mt_ (Tsuboi et al., 2009). Human RRF_mt_ is about 25-30 % identical to its eubacterial homologs but carries an additional 79 amino acids (aa) long extension at its N-terminus (Zhang and Spremulli, 1998). The recent high-resolution cryo-EM structures of RRF_mt_ bound to the 55S mitoribosomes (Koripella et al., 2019b) and an *in-vivo* formed mitoribosomal complex (Aibara et al., 2020) revealed that the structurally conserved segment of the RRF_mt_ is similar to its bacterial analog on (Agrawal et al., 2004; Barat et al., 2007; Dunkle et al., 2011; Fu et al., 2016; Gao et al., 2007; Gao et al., 2005; Weixlbaumer et al., 2007; Yokoyama et al., 2012; Zhou et al., 2020) and off (Kim et al., 2000; Nakano et al., 2003; Saikrishnan et al., 2005; Selmer et al., 1999; Toyoda et al., 2000; Yoshida et al., 2001) the 70S ribosome in terms of its overall size and domain composition. However, the unique mito-specific N-terminal extension (NTE) in RRF_mt_ extends towards the GTPase-associated center and interacts with the functionally important 16S rRNA elements of the mitoribosomal 39S subunit, including the rRNA helices 89 (H89), H90 and H92 (Koripella et al., 2019b).

Valuable mechanistic inferences about the bacterial ribosome recycling process were made from the structures of the 70S•RRF (e.g., Agrawal et al., 2004) and dissociated 50S•RRF•EF-G (Gao et al., 2007; Gao et al., 2005) complexes. Capturing the simultaneous binding of both factors on the 70S ribosome is challenging, however, owing to the rapid rate of 70S ribosomes dissociation into subunits by the combined action of RRF and EF-G (Borg et al., 2016). To slow down this reaction, a heterologous system with *T. thermophilus* RRF and *E. coli* EF-G was used to capture both factors on the 70S ribosome by cryo-EM (Yokoyama et al., 2012). Subsequently, a time-resolved cryo-EM study was also able to capture various 70S•RRF•EF-G functional intermediates, albeit at low resolution (Fu et al., 2016). More recently, a bacterial ribosome recycling complex containing both RRF and EF-G was obtained by X-ray crystallography by stabilizing EF-G on the 70S ribosome through a fusion between EF-G and ribosomal protein bL9 (Zhou et al., 2020). All these structures conclude that binding of EF-G to the 70S•RRF complex induces rotation of RRF domain II towards the helix 44 (h44) region of the 30S subunit, destabilizing the crucial intersubunit bridges B2a and B3, and thereby facilitating the dissociation of the 70S ribosome into its two subunits.

With a molecular weight of 87 kD, human EF-G2_mt_ is slightly larger than EF-G1_mt_ (83 kD), as well as both the isoforms of bacterial EF-G (78 kD) and EF-G2 (73 kD). It should be noted that some bacterial species do carry two isoforms of EF-G, but the function for the second bacterial isoform remains undefined (Connell et al., 2007; Margus et al., 2007; Seshadri et al., 2009), except in case of a spirochaete (Suematsu et al., 2010). EF-G2_mt_ has about 36 % aa sequence identity to EF-G1_mt_ and about 30 % aa identity to both its bacterial homologues. Some mammalian mitochondrial translation steps are now better understood through determination of the cryo-EM structures of the initiation (Koripella et al., 2019a; Kummer et al., 2018; Yassin et al., 2011) and the elongation (Koripella et al., 2020; Kummer and Ban, 2020) complexes at high-resolution. Our previous study of the human mitoribosome recycling complex of the RRF_mt_-bound 55S (Koripella et al., 2019b) provided useful insights into the mito-specific aspects of the recycling process, but a complete 55S mitoribosomal recycling complex comprising both RRF_mt_ and EF-G_mt_ remained elusive. To investigate the concerted action of RRF_mt_ and EF-G2_mt_ in splitting the 55S mitoribosome and to understand the detailed roles of mito-specific aa segments of RRF_mt_ and EF-G2_mt_ at the molecular level, we determined the key intermediate state structures of the 55S and 39S mitochondrial recycling complexes containing both RRF_mt_ and EF-G2_mt_.

## Results and discussion

### Structure of the human mitoribosome recycling complex

To investigate the molecular mechanism of ribosome recycling in mammalian mitochondria, we first prepared a model post-termination complex (PoTC) by incubating the human 55S mitoribosome with puromycin (Hirokawa et al., 2002; Koripella et al., 2019b). The model PoTC was briefly incubated with human RRF_mt_ and human EF-G2_mt_-GMPPCP to obtain the mitoribosome recycling complex. (see Supplemental Materials and Methods). Single-particle cryo-EM analysis on this complex yielded three major classes that each represent a major functional state formed during human mitoribosome recycling, referred henceforth to as Class I, Class II and Class III (Fig. S1). Class I corresponds to the intact 55S mitoribosome that carries a strong density for RRF_mt_, and was refined to 3.5 Å (Fig. 1A,D, S2A). Class II, a relatively small class with only 28,929 particle images, corresponds to the 55S mitoribosome that carries both RRF_mt_ and EF-G2_mt_, and was refined to 3.9 Å (Fig. S2B). In this class, the densities corresponding to the large (39S) mitoribosomal subunit, RRF_mt_, and EF-G2_mt_ are well resolved, but the small (28S) mitoribosomal subunit appears to be loosely bound and present in multiple poses (Fig. 1B, S2). The most populated Class III corresponds to the dissociated 39S subunits that carry both RRF_mt_ and EF-G2_mt_, and was refined to 3.15 Å (Fig. 1C,E, S2E). Class II likely represents an ensemble low-population intermediate states of mitoribosome recycling that occur between the states represented by Classes I and III. In addition to these three recycling complexes, we have also obtained a class of particles consisting of 55S mitoribosomes without either of the two factors where the 28S subunit was rotated by about 8° around its long axis such that its shoulder side moves closer to the 39S subunit while its platform side moves away from it (Fig. S3A). A similar orientation for the small subunit relative to the large subunit, termed as “subunit rolling” has been reported earlier for the 80S ribosomes (Budkevich et al., 2014) and the 55S mitoribosomes (Amunts et al., 2015; Koripella et al., 2019b).

**Fig. 1.**
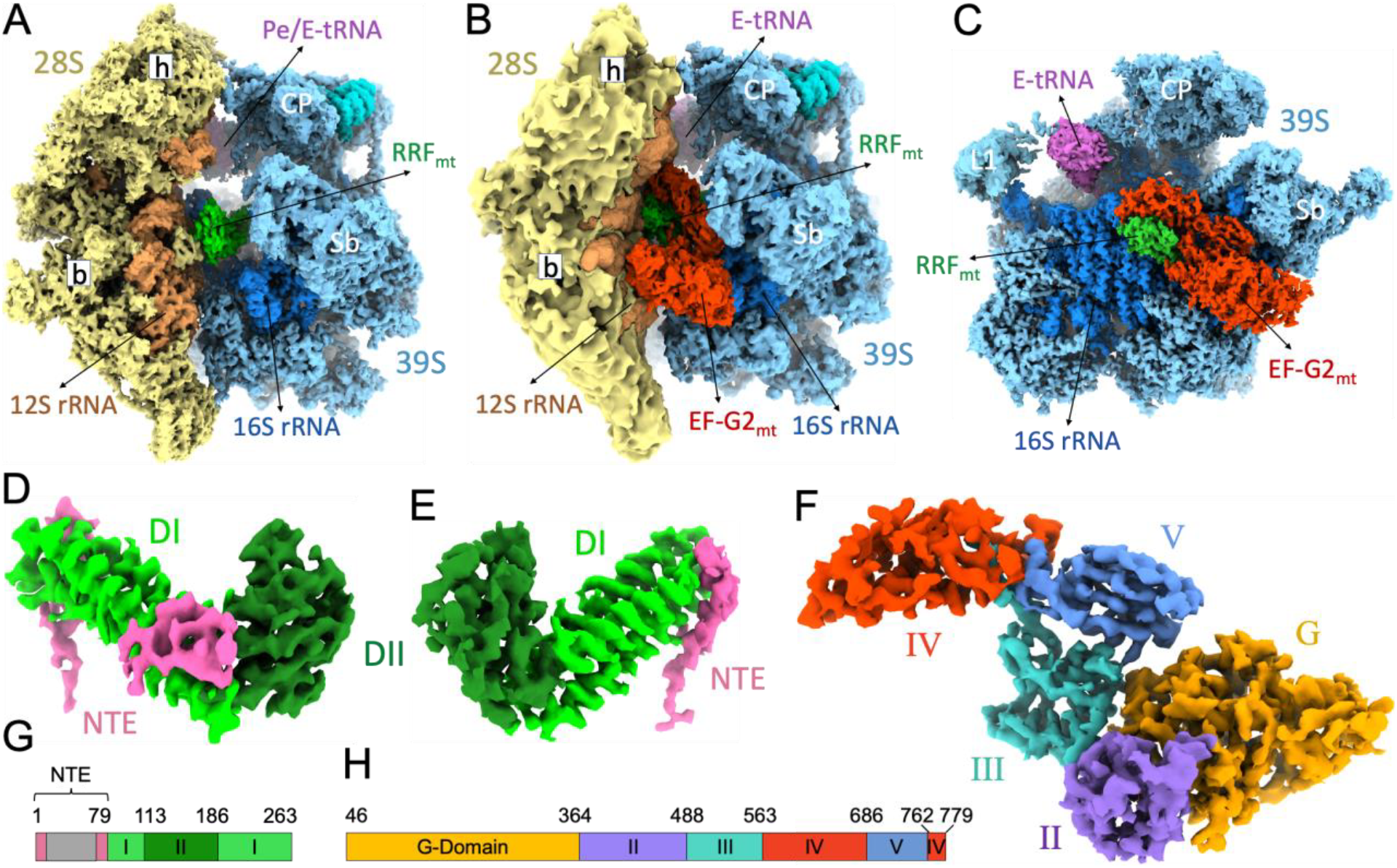
Cryo-EM structures of the human mitochondrial recycling complexes in three functional states. Segmented cryo-EM maps of the mitoribosomal **(A)** 55S•RRF_mt_ complex (Class I), **(B)** 55S•RRF_mt_•EF-G2_mt_•GMPPCP complex (Class II), and **(C)** 39S•RRF_mt_•EF-G2_mt_•GMPPCP complex (Class III). In these three panels, the 28S subunit is shown in yellow, the 39S subunit in blue, E-tRNA in orchid, RRF_mt_ in green and EF-G2_mt_ in red. A lighter shade of yellow differentiates the 28S ribosomal proteins from the 12S rRNA, while a lighter shade of blue differentiates the 39S ribosomal proteins from the 16S rRNA. Landmarks of the 28S subunit: h, head; b, body. Landmarks of the 39S subunit: CP, central protuberance; Sb, stalk base; L1, MRP uL1m. **(D)** Cryo-EM density of RRF_mt_ extracted from the Class I complex. **(E)** Cryo-EM density of RRF_mt_ extracted from the Class III complex. **(F)** Cryo-EM density of EF-G2_mt_ extracted from Class III complex. See Figs. S1 and S2, for overall and local resolutions, respectively, of densities corresponding to the mitoribosome, RRF_mt_, and EF-G2_mt_ in these complexes. **(G)** Domain organization in RRF_mt_ displaying domain I (light green), domain II (dark green), modelled region of NTE (pink) and unmodelled region of NTE (gray) **(H)** Domain organization in EF-G2_mt_ showing G domain (orange), domain II (purple), domain III (cyan), domain IV (red) and domain V (blue).

### RRF_mt_ binding stabilizes the rotated state of the 28S subunit in the 55S mitoribosome

Superimposition of our Class I complex with the RRF_mt_-unbound human (Amunts et al., 2015), bovine (Sharma et al., 2003) and porcine (Greber et al., 2015) 55S mitoribosomes showed that the small 28S subunit was rotated counter-clockwise by about 8.5° with respect to the large 39S subunit (Fig. S3B), similar to the “ratchet-like inter-subunit rotation” observed in the bacterial 70S ribosome (Frank and Agrawal, 2000, 2001) and the 55S mitoribosomes complexed with translational factors (Koripella et al., 2020; Koripella et al., 2019b; Kummer and Ban, 2020; Kummer et al., 2018). In addition to the inter-subunit rotation, the head domain of the 28S subunit is rotated by about 4° towards the tRNA exit (E) site in a direction roughly orthogonal to the inter-subunit motion (Fig. S3B), similar to “head swiveling” in the bacterial 70S ribosomes (Ratje et al., 2010; Schuwirth et al., 2005). As expected, the structure of the Class I complex matches the previously published 3.9 Å resolution map of the analogous 55S•RRF_mt_ complex (Koripella et al., 2019b).

The Class I map showed the characteristic “L” shaped RRF_mt_ density and a density corresponding to a pe/E-state tRNA within the inter-mitoribosomal subunit space. The overall positioning and domain arrangement of RRF_mt_ in the Class I map is similar to the bacterial RRF on (Agrawal et al., 2004; Barat et al., 2007; Dunkle et al., 2011; Fu et al., 2016; Gao et al., 2007; Gao et al., 2005; Weixlbaumer et al., 2007; Yokoyama et al., 2012; Zhou et al., 2020) and off (Kim et al., 2000; Nakano et al., 2003; Saikrishnan et al., 2005; Selmer et al., 1999; Toyoda et al., 2000; Yoshida et al., 2001) the 70S ribosomes, and also to the structures of RRF bound to the 70S chloroplast ribosome (Boerema et al., 2018; Sharma et al., 2007), and RRF_mt_ bound to the human mitochondrial 55S in our previous study (Koripella et al., 2019b). As observed in bacteria, domain I is positioned close to the peptidyl transferase center (PTC) and extends towards the α-sarcin-ricin loop (SRL). A striking difference between the human RRF_mt_ and its bacterial counterpart is the presence of a 79 aa long N-terminal extension (NTE) in RRF_mt_. We could model the last 14 aa residues of the NTE of RRF_mt_ into an additional density contiguous with the α-helix1 from domain I. As discussed in our previous study (Koripella et al., 2019b), the NTE is strategically positioned in the intersubunit space between domain I and several functionally important 16S rRNA structural elements such as H89, H90, H92 (A-loop) and MRP L16 (Fig. S4) and interacts with several nucleotides (nts) and aa residues in its vicinity (Koripella et al., 2019b). Interestingly, unlike the α-helical nature inferred for the part of this segment of NTE (Koripella et al., 2019b), we find that its higher resolution density to be partially unstructured. A similar observation of a relatively unstructured NTE has been recently reported in an *in-vivo* state complex (Aibara et al., 2020). However, the mitoribosomal components interacting with RRF_mt_ as described before (Koripella et al., 2019b) essentially remain unaltered.

We found a small density in a tight pocket surrounded by the outer bend of the junction between domains I and II of RRF_mt_, MRP uS12m and the small subunit’s 12S rRNA helix h44, and the large subunit’s 16S rRNA helices H69 and H71 (Fig. 2A). Except for our previous lower resolution map of the 55S•RRF_mt_ complex (Koripella et al., 2019b), this additional density is not observed in any of the available 55S mitoribosomal structures, whether complexed with other translational factors (Koripella et al., 2020; Kummer and Ban, 2020; Kummer et al., 2018) or not (Amunts et al., 2015; Greber et al., 2015). Since our complex was reconstituted from purified components, this additional density should correspond to an RRF_mt_ NTE segment that has been stabilized through interactions with multiple mitoribosomal components in its vicinity. Though bacterial and mitochondrial ribosomes exhibit significant differences in their overall shape, composition, and conformation, their internal rRNA core regions are largely conserved (Amunts et al., 2015; Greber et al., 2015; Kaushal et al., 2014; Sharma et al., 2003). Comparison of the RRF binding sites between the bacterial and mitochondrial ribosomes reveals that the H69 of 16S rRNA is slightly shorter in the mammalian mitoribosomes (Fig. 2B). This minor shortening of H69 is critical because it directly impacts the interaction of H69 with domain II of RRF_mt_. The small density most likely corresponding to N-terminus segment of the mito-specific NTE appears to compensate for the shortened H69 by mediating the interactions between RRF_mt_ and H69 (Fig. 2B). Based on the available sidechain information from the small density, we generated a model that would account for the 10 aa residues (Ala2-Val11) at the N-terminus of NTE (Fig 2B). The present model is guided primarily by density for two consecutive large side chains of Phe8 and Arg9 within the first 11 aa (Fig. 2C). Since at least three more combination of two consecutive aa with large sidechains are present in the unmodelled segment of NTE, we refrain from analyzing the molecular interactions of this small segment of NTE with the rest of the complex.

**Fig. 2.**
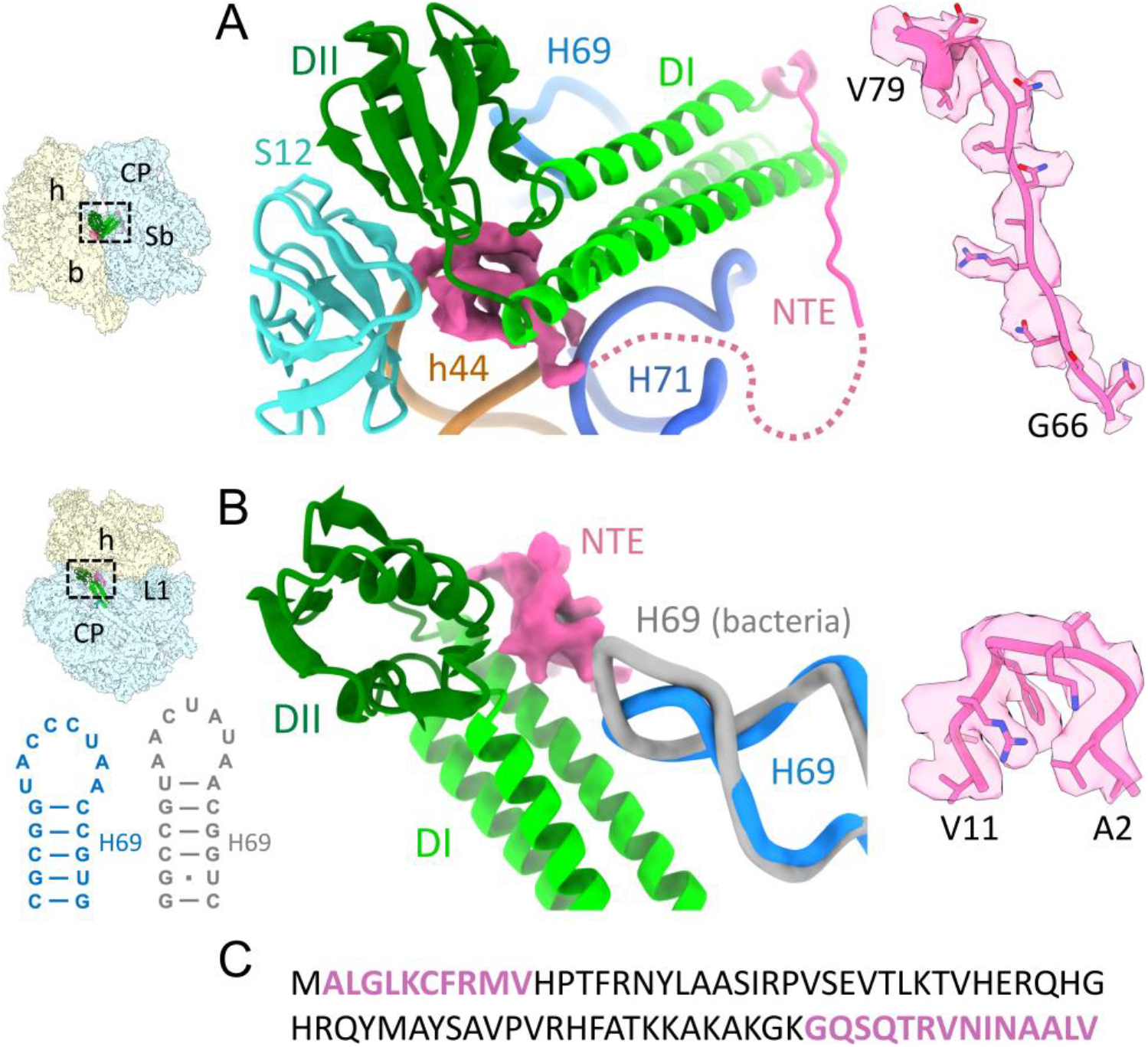
Structures of two NTE segments of RRF_mt_. **(A)** Density corresponding to the N-terminus segment of RRF_mt_ NTE (pink) observed in the inter-subunit space sandwiched between the 28S SSU components, including MRP uS12m (cyan) and the 12S rRNA helix h44 (brown), the 39S LSU components, including 16S rRNA helices H69 (medium blue) and H71 (dark blue), and the outer bent of the junction between domains I and II of RRF_mt_. The dotted line (pink) depicts the connection between two structurally stabilized segments of the RRF_mt_‘s NTE. **(B)** H69 superimposition from bacterial (gray) (Weixlbaumer et al., 2007) and human mitochondrial ribosomes (blue) reveals the shortening of H69 in the mitoribosome, also depicted in secondary structures of H69 on the lower left. Domain II of RRF_mt_ interacts with the shorter H69 through its strategically positioned N-terminus of NTE (pink). **(C)** Sequence of NTE showing two modeled regions (pink). Thumbnails to the left depicts an overall orientation of the 55S mitoribosome, with semitransparent 28S (yellow) and 39S (blue) subunits, and overlaid positions of ligands. Landmarks on the thumbnail: h, head, and b, body of the 28S subunit, and CP, central protuberance; Sb, stalk base of the 39S subunit.

It should be noted that the RRF_mt_-bound 55S mitoribosomes (present work and (Koripella et al., 2019b)) were never observed in the unrotated state, suggesting that the RRF_mt_ binding locks the ribosome in a fully rotated state. This is in contrast to the bacterial 70S•RRF complexes that were found both in their rotated and unrotated conformational states (Agrawal et al., 2004; Barat et al., 2007; Dunkle et al., 2011; Fu et al., 2016; Gao et al., 2007; Gao et al., 2005; Weixlbaumer et al., 2007; Yokoyama et al., 2012; Zhou et al., 2020). The rotated conformation of the 55S mitoribosome seems to prime subunit dissociation by either destabilizing or completely breaking seven out of fifteen inter-subunit bridges in the unrotated 55S mitoribosome (Amunts et al., 2015; Sharma et al., 2003). The simultaneous interactions of N-terminus segment of the RRF_mt_ NTE with RRF_mt_’s structurally conserved domain I, MRP uS12m, h44, H69 and H71 likely help prevent the back-rotation of the small 28S subunit. The rotated state of the 55S mitoribosome could serve as an ideal substrate for the subsequent binding of EF-G2_mt_ to complete subunit dissociation. In this context, it is important to note that the entire 79 aa long NTE is an integral part of the mature protein and is known to be essential for RRF_mt_ function during the mitoribosome recycling (Rorbach et al., 2008; Zhang and Spremulli, 1998).

### RRF_mt_ domain II motion helps split the 55S mitoribosome into its two subunits

Using fast-kinetics, it has been shown that the splitting/recycling of the 70S ribosome by the concerted action of RRF and EF-G happens in the sub-second time scale (Borg et al., 2016). Both RRF and EF-G have been captured on the 70S ribosome with time-resolved cryo-EM (Fu et al., 2016). It is more challenging without time-resolved techniques, but RRF and EF-G were also captured on the 70S ribosomes by using the factors from different species (Yokoyama et al., 2012) or by crosslinking the EF-G with one of the ribosomal proteins (Zhou et al., 2020). In the present work, collection of very large cryo-EM datasets (altogether 21,752 micrographs, (Fig. S1) enabled the isolation of a small subset of 55S particles (Class II) that contained both RRF_mt_ and EF-G2_mt_ (Fig. 1B). However, the 28S subunit density was found to be weak and present in multiple destabilized conformations relative to the 39S subunit in this complex.

Both Class II and Class III maps showed readily recognizable densities corresponding to RRF_mt_ and EF-G2_mt_. The Class III complex had superior resolution, which enabled more accurate analysis of molecular-level interactions between the two factors and the mitoribosome, and their functional implications, while the Class II map was useful for interpreting large-scale conformational changes. In line with the bacterial 70S/50S•RRF•EFG complexes (Fu et al., 2016; Gao et al., 2007; Gao et al., 2005; Yokoyama et al., 2012; Zhou et al., 2020), the conformation of RRF_mt_ is substantially different between the Class I and Class III recycling complexes. The conformation of RRF_mt_ domain I remains unchanged among all three classes. In the Class II and III maps, domain II was rotated by about 45° towards the small subunit compared to its position in the Class I complex (Fig. 3A). This large conformational change is enabled by a highly flexible hinge regions between domain I and domain II in RRF_mt_. Due to this rotation, the tip of domain II moved by about 40 Å towards the h44 of 12S rRNA. When the maps of Class II and III were superimposed, this motion resulted in a major steric clash between the RRF_mt_ domain II and the 28S subunit elements h44 and MRP uS12m (Fig. 3A). In the 55S mitoribosome, h44 is involved in the formation of two intersubunit bridges B2a and B3 by pairing with H69 and H71, respectively (Amunts et al., 2015; Sharma et al., 2003). By displacing h44, RRF_mt_ disrupts these crucial intersubunit bridges, thereby splitting the 55S mitoribosome into its two subunits.

**Fig. 3.**
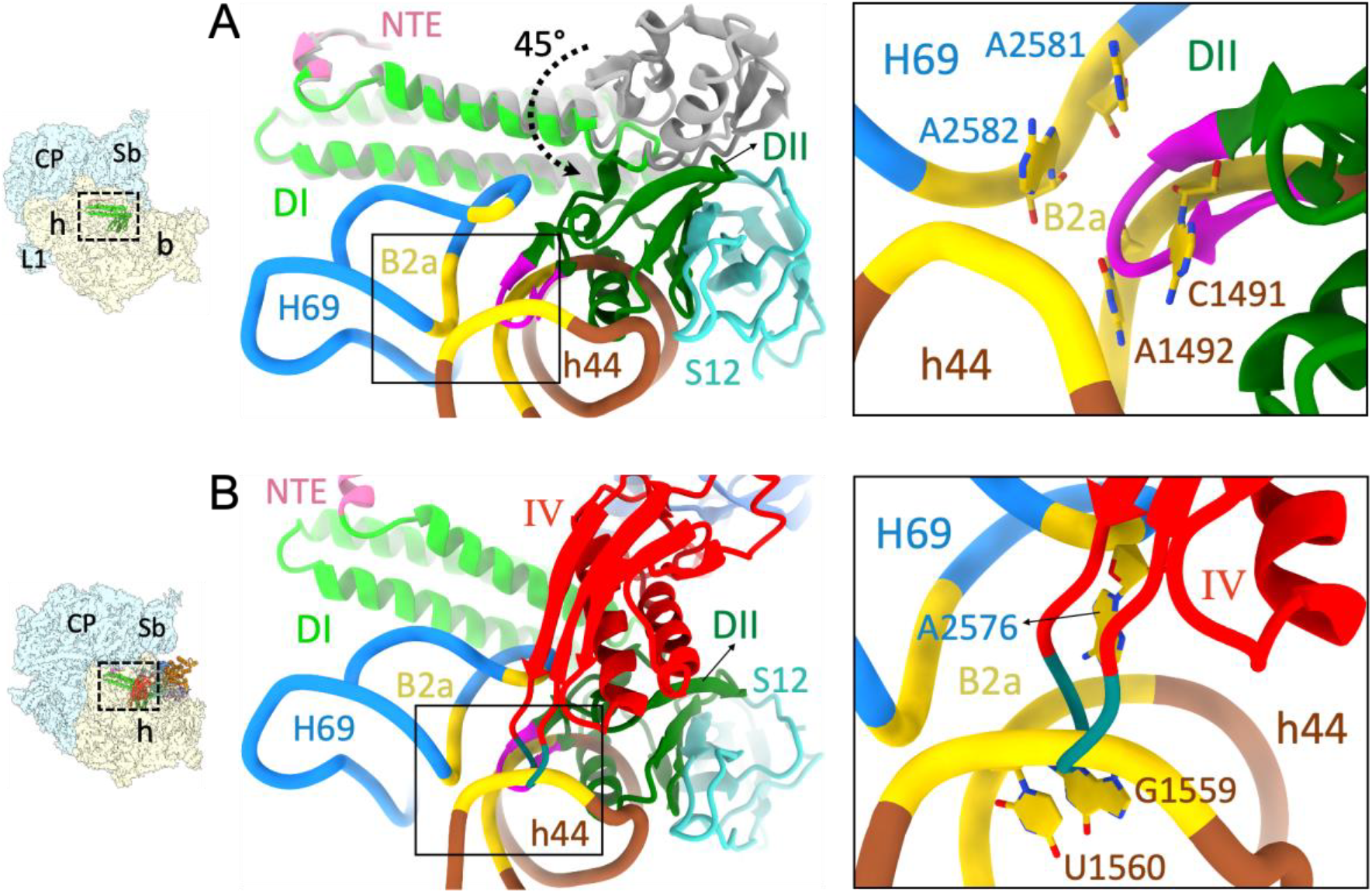
Direct involvement of RRF_mt_ domain II and EF-G1_mt_ domain IV in human mitoribosome recycling. **(A)** Comparison of the overall conformation of RRF_mt_ between the Class I (gray) and Class III (green) complexes revealed that domain II rotates by about 45° towards the 28S subunit. Such a rotation would sterically clash with the 12S rRNA helix, h44 (brown) and MRP uS12m (cyan) and destabilize a crucial inter-subunit bridge, B2a (yellow). Inset shows the magnified view of the inter-subunit bridge B2a with RRF_mt_ domain II residues V121-K127 (red) disrupting B2a by inserting between h44 residues (C1491 and A1492) and H69 residues (A2581 and A2582). **(B)** Superimposition of the Class III complex with the Class I complex shows a direct overlap between the loop1 region (dark cyan) of EF-G2_mt_ domain IV (red) and the intersubunit bridge B2a (yellow) formed between h44 (brown) and H69 (blue). Inset shows the magnified view of EF-G2_mt_ domain IV residues L582-R585 (dark cyan) that will disrupt the bridge B2a by inserting between h44 residues (G1559 and U1560) and H69 residue (A2576) and pushing the 28S subunit (h44) away. Thumbnails to the left depict an overall orientation of the 55S mitoribosome, with semitransparent 28S (yellow) and 39S (blue) subunits, and overlaid position RRF_mt_. Landmarks on the thumbnails are same as in Fig. 2.

A well-defined 28S structure was not seen within the Class II 55S mitoribosomal complex, but several inter-subunit bridges do appear to be destabilized or broken, and the small subunit seems to be dissociating from the large subunit. To remove any possible large subunit contamination from the Class II map, extensive reference-based 3D classification was employed, but this class of 55S mitoribosomal particles with a well-resolved 39S subunit and a poorly resolved 28S subunit remained unchanged. This supports it being an ensemble of authentic, short-lived functional intermediates of mitoribosome recycling, where the small subunit is captured in multiple positions during its separation from the large subunit.

### EF-G2_mt_ binding induces conformational changes in both RRF_mt_ and the mitoribosome

The 55S•RRF_mt_ complex (Class I) undergoes large conformational changes upon binding to EF-G2_mt_ (Classes II and III). EF-G2_mt_ would not be able to access its binding site on the mitoribosome without the significant movement seen in domain II of RRF_mt_ in Class II and Class III complexes. This movement eliminates the direct spatial conflicts of EF-G2_mt_ domains III, IV and V with the initial position of RRF_mt_ domain II in the Class I complex (Fig. 3A,B; Fig. S5). This change could be induced either during or upon binding of EF-G2_mt_. The domain II of RRF_mt_ is repositioned into a cavity created by domains III, IV and V of EF-G2_mt_ (Fig. S6A) and an extensive network of interactions are formed between the two mitochondrial recycling factors. This is also in agreement with the location of RRF domain II reported in the bacterial 50S (Gao et al., 2007) and 70S complexes (Fu et al., 2016; Yokoyama et al., 2012). A majority of the interactions are formed between the hinge regions that connect the two domains of RRF_mt_ and the loop regions from domain III of EF-G2_mt_. Several aa residues from the RRF_mt_ hinge region (Pro183-Thr186) interact with EF-G2_mt_ domain III residues Glu495-Leu499 (Fig. S6B) via hydrogen bonds and hydrophobic interactions. Arg187 from the α-helix following the hinge region of RRF_mt_ domain II has close hydrogen-bonding interactions with Tyr556 of EF-G2_mt_ domain III (Fig. S6B). A second set of contacts formed between the two factors involves residues Ile109-Arg110 from the second hinge region of RRF_mt_ and residues Ser527-Gln529 from the domain III of EF-G2_mt_ (Fig. S6C).

Domain IV of EF-G2_mt_ presses against domain II of RRF_mt_ through multiple interactions. The surface residues of α-helix 1 (Asn629, Ser633 and Leu636) and α-helix 2 (Thr664, Met665, Ser667 and Ala668) from domain IV of EF-G2_mt_ interact with residues in β-strand 3 (Ser134-Met138) and its adjoining loop region (Gln132 and Ile133) from domain II of RRF_mt_ through a combination of electrostatic and hydrophobic interactions (Fig. S6D). Gln637 from the α-helix 1 of EF-G2_mt_ also shares a hydrogen bond with Ser112 from the hinge region of RRF_mt_ (Fig. S6D). Contacts are observed between the C-terminal α-helix of EF-G2_mt_ domain IV and the α-helix 3 from the triple-helix bundle of RRF_mt_ domain I. In bacteria, the analogous C-terminal α-helix of EF-G is often considered as part of domain V though the first atomic models of EF-G (AEvarsson et al., 1994; Czworkowski et al., 1994) grouped it with domain IV. While residues Ser776-Leu778 from EF-G2_mt_ domain IV pair with residues Arg251 and Val255 from RRF_mt_ domain I through hydrogen bonds (Fig. S6E), Arg775 from EF-G2_mt_ domain IV strongly interacts with Glu259 from RRF_mt_ domain I through a salt-bridge (Fig. S6E). Direct interactions of EF-G2_mt_ domain III with RRF_mt_ domain II at its hinge regions, known to confer interdomain flexibility to the bacterial factor (Nakano et al., 2003; Saikrishnan et al., 2005; Yoshida et al., 2001), likely help trigger the dissociation of 55S mitoribosomes into subunits by enabling the repositioning of RRF_mt_ domain II. At the same time, the multiple interactions between domain IV of EF-G2_mt_ and domain II of RRF_mt_ appear to stabilize the RRF_mt_ domain II in the altered position, which would push the 28S subunit away from the 39S subunit and prevent domain II from reverting back to its previous orientation, in order to maintain the 39S•RRF_mt_•EF-G2_mt_ complex in a dissociated state.

In addition to aiding to the function of RRF_mt_, EF-G2_mt_ plays a direct role in destabilizing the 55S mitoribosome. Superimposition of the maps of Class I and Class III complexes reveals a direct steric clash between the loop1 region of EF-G2_mt_ domain IV and the 28S subunit component (rRNA h44) that participates in the formation of the inter-subunit bridge B2a (Fig. 3B). More importantly, the orientation of domain IV loop1 seems to be unique to EF-G2_mt_ since the analogous region in EF-G1_mt_ is positioned away from the intersubunit bridge B2a towards the decoding center (Koripella et al., 2020; Kummer and Ban, 2020).

EF-G2_mt_ binding resulted in a prominent conformational change in the uL11m stalk-base region of the 39S subunit. The uL11m stalk-base region moved towards the CTD of uL12m, a component of the L10-L12 stalk, and assumed a unique conformation not reported previously (Amunts A 2015, Greber BJ 2015, Kummer E 2018, Koripella RK 2019; 2020). The 16S rRNA H43 of the uL11m stalk-base moved by 3 Å towards the CTD of uL12m, oriented parallel to the domain V of EF-G2_mt_, while the NTD of MRP uL11m moved about 5 Å away from the EF-G2_mt_ domain V (Fig. 4A). This is in sharp contrast to the 55S•EF-G1_mt_ translocation complexes (Amunts et al., 2015; Greber et al., 2015; Koripella et al., 2020; Koripella et al., 2019b; Kummer and Ban, 2020; Kummer et al., 2018), where the uL11m stalk-base region was observed to move 5 Å closer towards the domain V of EF-G1_mt_ (Fig. 4B). The movement of uL11m away from the EF-G2_mt_ domain V and towards the CTD of uL12m is essential for the binding of EF-G2_mt_ in the present conformation, to avoid the steric clash between the NTD of uL11m and domain V of EF-G_mt_ (Fig. 4A). It is also possible that the conformation of EF-G2_mt_ observed in our 39S•EF-G2_mt_ complex (Class III) was attained after the dissociation of 39S subunit from the 55S complex (Class II).

**Fig. 4.**
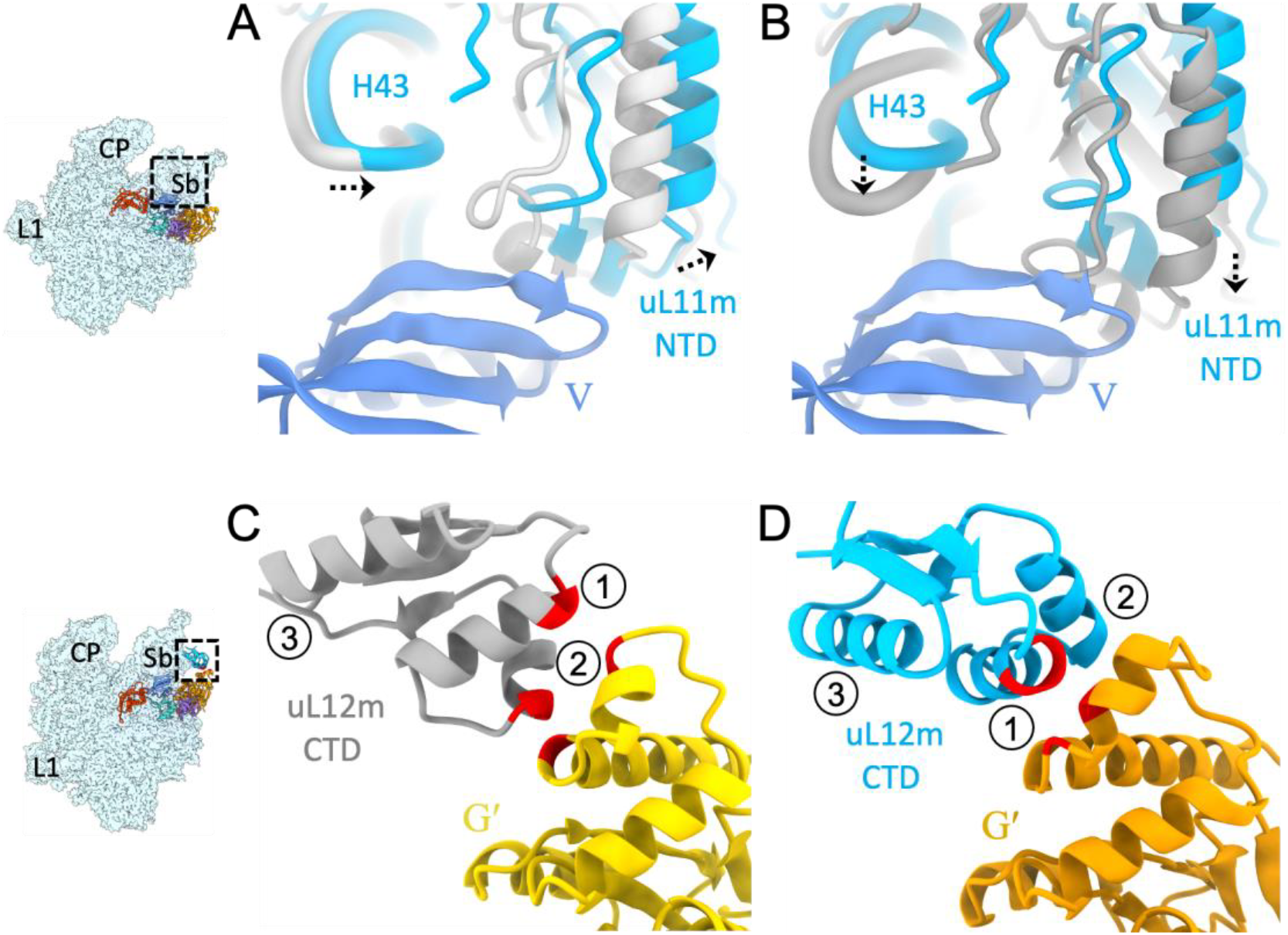
EF-G2_mt_ binding induces large-scale conformational changes in the uL11m stalk-base region and the CTD of uL12m. **(A)** In the 39S•RRF_mt_•EF-G2_mt_ complex, the uL11m stalk-base region (blue) moves towards the CTD of uL12m and away from the domain V of EF-G2_mt_, as compared to its position in the EF-G2_mt_-unbound 55S mitoribosome (light gray) (Amunts et al., 2015). **(B)** In sharp contrast, in the presence of EF-G1_mt_, the uL11m stalk-base region was found to move towards the domain V of EF-G1_mt_ in the translocation complex (dark gray) (Koripella et al., 2020). **(C)** In the 55S•EF-G1_mt_ complex (Koripella et al., 2020), the CTD of uL12m (gray) is positioned in such a way that its α-helices 1 and 2 directly interact with the G′ subdomain of EF-G1_mt_ (yellow). **(D)** In our 39S•RRF_mt_•EF-G2_mt_ complex, the CTD of uL12m (blue) is rotated by about 60° and shifted away by about 7 Å from the G′ subdomain of EF-G2_mt_, resulting in the loss of contacts between its α-helix 2 and G′ subdomain of EF-G2_mt_ while a new set of interactions is formed between its α-helix 1 and G′ subdomain of EF-G2_mt_. Thumbnails to the left depict an overall orientation of the 39S subunit (semitransparent blue), and overlaid positions of the ligands. Landmarks on the thumbnail: CP, central protuberance; Sb, stalk base; L1, MRP uL1m.

In addition to the unique conformation of the uL11m stalk-base region, the CTD of uL12m was also observed in a distinct conformation. In the 55S•EF-G1_mt_ complexes (Koripella et al., 2020; Kummer and Ban, 2020), the CTD of uL12m is positioned close to EF-G1_mt_ so that α-helices 1 and 2 of uL12m CTD would have close interactions with the G′ subdomain of EF-G1mt (Fig. 4C), thereby providing stability to the otherwise flexible uL12m CTD. In contrast, the CTD of uL12m rotates by about 60° and shifts away by about 7 Å from the G′ subdomain of EF-G2_mt_ in the 39S•EF-G2_mt_ complex (Fig. 4D). As a result of this large rotational movement, interactions between the uL12m CTD α-helix 2 and the G′ subdomain of EF-G2_mt_ are lost, while the contacts between the α-helix 1 and the G′ subdomain of EF-G2_mt_ are maintained (Fig. 4D). Since protein uL12 is known to play a central role in the recruitment of translational factors to the bacterial ribosome (Datta et al., 2005; Diaconu et al., 2005; Helgstrand et al., 2007; Imai et al., 2020) the semi-stable conformation of uL12m observed in the 39S•EF-G2_mt_ complex represents a late-stage conformation of uL12m prior to its detachment from the EF-G2_mt_ as EF-G2_mt_’s participation in the 55S ribosome recycling process nears completion.

### Structural basis for use of EF-G2_mt_ in mitoribosomal recycling instead of EF-G1_mt_

In most bacterial species, a single EF-G is involved in both translocation and ribosome recycling. Mammalian mitoribosomes have evolved to utilize the two isoforms EF-G1_mt_ and EF-G2_mt_ to cope with the significantly altered environment in mitochondria as compared to the bacterial cytoplasm (Hammarsund et al., 2001; Tsuboi et al., 2009) and perform two separate functions (Hammarsund et al., 2001; Tsuboi et al., 2009). EF-G1_mt_ is used during translation elongation while EF-G2_mt_ is used along with RRF_mt_ for the recycling of the 55S mitoribosome. The overall position and domain arrangement of EF-G2_mt_ is similar to that of EF-G1_mt_ in the recently published human (Koripella et al., 2020) and porcine (Kummer and Ban, 2020) translocational complexes (Fig. S5). There are, however, some specific structural distinctions between the two factors that assign them specialized functional roles. The most striking of which is the presence of a C-terminal extension (CTE) in EF-G1_mt_ domain IV (Fig. 5A) (Koripella et al., 2020). Besides its CTE, the size of the conserved C-terminal α-helix of EF-G1_mt_ domain IV is substantially longer (16 aa) (Fig. 5A) than the C-terminal α-helix (12 aa) of EF-G2_mt_ domain IV (Fig. 5B).

**Fig. 5.**
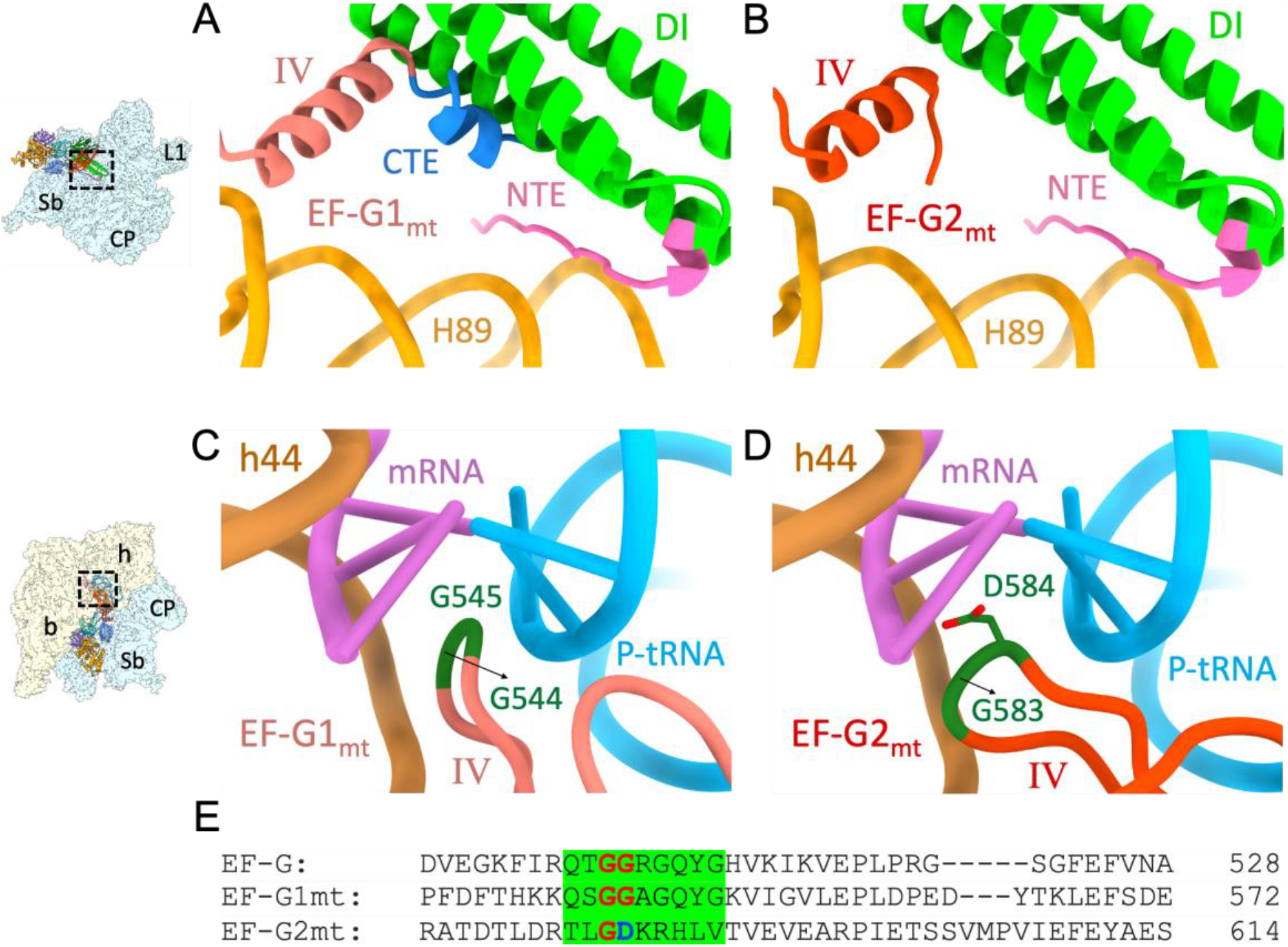
Structural basis for the exclusive roles of EF-G1_mt_ in tRNA translocation and EF-G2_mt_ in mitoribosome recycling. **(A)** The presence of a substantially longer C-terminal α-helix (salmon) along with the presence of a unique CTE (blue), which is required for mitochondria tRNA translocation (Koripella et al., 2020), prevents the simultaneous binding of EF-G1_mt_ and RRF_mt_ (green) on the mitoribosome due to a major steric clash. Furthermore, any conformational repositioning the C-terminal α-helix of EF-G1_mt_ will result in direct steric clash with the H89 (orange) of 16S rRNA and the NTE of RRF_mt_ (pink)**. (B)** Having a shorter C-terminal α-helix (red) and the absence of CTE allows the simultaneous binding of EF-G2_mt_ and RRF_mt_ (green) on the mitoribosome. **(C)** In EF-G1_mt_ (Koripella et al., 2020), the presence of two universally conserved glycine residues (dark green) at the tip of domain IV loop1 region (salmon) facilitates its insertion between the mRNA-tRNA duplex at the decoding center (DC) during EF-G1_mt_ - catalyzed translocation. **(D)** In EF-G2_mt_, the second glycine of domain IV loop 1 region (red) is substituted by an aspartic acid (dark green) altering its conformation and flexibility, and thereby making its insertion into the DC during translocation unfavorable. **(E)** The aa sequence highlighted in green corresponds to loop 1 situated at the tip of domain IV, which is highly conserved between EF-G1_mt_ and the *T. thermophilus* EF-G. The universally conserved glycine residues are shown in red while the aspartic acid substitution in EF-G2_mt_ is shown in blue.

The sturdier and longer C-terminal α-helix of EF-G1_mt_ domain IV and its 11 aa CTE would not permit the coexistence of RRF_mt_ on the mitoribosome due to a major steric clash between domain I of RRF_mt_ and the C-terminal region of EF-G1_mt_ (Fig. 5B). Even a reorientation of the C-terminal α-helix and its CTE away from the domain I of RRF_mt_ would not resolve the problem as they would then clash with H89 of the 16S rRNA and the NTE of RRF_mt_ that has been positioned in the inter subunit space between the domain I of RRF_mt_ and 16S rRNA helix, H89 (Fig. 5B). Structural analysis of the bacterial EF-Gs from various species (Chen et al., 2010; Gao et al., 2009; Pulk and Cate, 2013) has revealed that the length of their C-terminal α-helices are about 12 aa long, suggesting that only EF-G2_mt_ can function alongside RRF_mt_ during subunit splitting. Moreover, the interaction between the C-terminal regions of EF-G2_mt_ domain IV and RRF_mt_ domain I is essential for stabilizing the bound RRF_mt_. This tight anchoring of RRF_mt_ domain I to the mitoribosome would prevent RRF_mt_ dissociation from the mitoribosome when its domain II undergoes substantial rotation to displace the h44 region of the 28S subunit. This agrees with the observation that deletion of the last few aa residues from the C-terminal region of bacterial RRF adversely effects its function during 70S ribosome recycling (Fujiwara et al., 2001; Fujiwara et al., 1999).

Now the question is why EF-G1_mt_ mediates the translocation step during mitoribosomal elongation and not EF-G2_mt_? The most probable answer is that the CTE in the domain IV of EF-G1mt that limits its ability to participate in the mitochondrial ribosome recycling, is reported to be directly involved in the elongation step of mammalian mitochondrial protein synthesis (Koripella et al., 2020). By contacting the inner bend of the A-site tRNA, the CTE of EF-G1_mt_ can help in translocating the CCA arm of the A-site tRNA into the P site, and by interacting with a 16S rRNA helix, H71, it prevents the P-site tRNA from reverting back into the A site (Koripella et al., 2020). This analysis is further supported by the fact that bacterial EF-Gs, which lack the CTE, are inactive on the 55S mitochondrial ribosomes while EF-G1_mt_ is active on the 70S ribosomes (Eberly et al., 1985).

Specific sequence differences in a critical region (loop1) within the domain IV of EF-G2_mt_ can also make it ineffective in driving translocation. During EF-G-catalyzed translocation in bacteria, the presence of two universally conserved glycine residues at the tip of domain IV (loop 1) region facilitate the insertion of domain IV into the decoding center (DC) (Gao et al., 2009). In the DC, the loop 1 of domain IV destabilizes the codon-anticodon interactions of the mRNA-tRNA duplex with the universally conserved 16S rRNA bases A1492 and A1493 in the 30S A site thereby aiding the A-site tRNA along with its associated codon to translocate into the P site (Zhou et al., 2014). The loop 1 region is conserved in EF-G1_mt_ but is significantly altered in EF-G2_mt_ (Fig. 5E). EF-G1_mt_ retains both the glycine residues (Gly544 and Gly545) in its domain IV loop 1 region (Fig. 5C) whereas the second glycine is replaced by an aspartic acid in EF-G2_mt_ (Fig. 5D) which alters the conformation of the tip of domain IV, rendering it structurally unfavorable for insertion into the grove between the P-site tRNA and the associated codon (Fig. 5D). Moreover, the Ala546 and Gly547 residues that follow the conserved glycines of loop 1 in EF-G1_mt_ are substituted by the large polar sidechain residues lysine and arginine, respectively, in the corresponding region of EF-G2_mt_ (Fig. 5E), thereby significantly altering the hydrophobicity of the loop 1 tip. Point mutations and deletions at the loop 1 tip region are known to have a pronounced effect on the function of EF-G during the elongation step of bacterial protein synthesis (Kolesnikov and Gudkov, 2003) but have a negligible effect on the activity of EF-G in bacterial ribosome disassembly (Zhang et al., 2015). Our study thus provides a structural rationale for the biochemical finding that mammalian mitoribosomes utilize two distinct EF-G-like factors during the translation elongation and recycling phases (Tsuboi et al., 2009).

### Mammalian 55S mitoribosome recycling does not require GTP hydrolysis by EF-G2_mt_

The overall G domain structure in the 39S•RRF_mt_•EF-G2_mt_ complex (Class III map) is similar to the G domain in the 55S•EF-G1_mt_ complex (Koripella et al., 2020; Kummer and Ban, 2020). The well-ordered density in our maps of highly conserved translational GTPase consensus motifs such as the P-loop, switch I and switch II regions, allowed complete modeling of these essential regions. A well-defined density corresponding to a bound GMPPCP molecule is also readily identifiable in the nucleotide binding pocket. GMPPCP is stabilized through interactions with universally conserved aa residues, such as Asp80 and Lys83 of P-loop, Thr122 of switch I and His145 of switch II (Fig. S7A). As in the 55S•EF-G1_mt_ complex (Koripella et al., 2020), a crucial Mg^2+^ ion is positioned near the γ phosphate of GMPPCP and is coordinated by Thr84 and Thr122 from the P-loop and switch I regions, respectively (Fig. S7A). The catalytic His145 (His124 in EF-G1_mt_), known to play a central role in the hydrolysis of the bound nucleotide (Chen et al., 2013; Tourigny et al., 2013), is found oriented towards the γ phosphate of the bound GMPPCP (Fig. S7A), suggesting an active conformation of the factor prior to GTP hydrolysis. The highly conserved α-sarcin-ricin stem-loop (SRL) region is known to be essential for the GTPase activity of all the translational G proteins (Chen et al., 2013; Gao et al., 2009; Tourigny et al., 2013). Base A3129 from the SRL was found to be contacting the Switch II His145 through hydrogen bonding interactions and thereby stabilizing His145 in its activated conformation poised to perform the hydrolysis reaction (Fig. S7B). While EF-G-dependent GTP hydrolysis is essential for an efficient splitting of the 70S ribosome into its subunits (Borg et al., 2016; Karimi et al., 1999; Zavialov et al., 2005), dissociation of the mammalian 55S mitoribosomes does not require EF-G2_mt_-dependent GTP hydrolysis, which is only needed for the release of EF-G2_mt_ from the dissociated large subunit (Tsuboi et al., 2009). The above hypothesis is strongly supported by our observation that a significant proportion (82 %) of the EF-G2_mt_ was found complexed to the 39S subunits as compared to the small proportion (18 %) that remained associated with the 55S mitoribosomes (see Fig. S1). Even though GTP hydrolysis by EF-G2_mt_ is not necessary for 55S mitoribosome splitting, the presence of GTP or its non-hydrolysable analogues GDPNP/GMPPCP in the nucleotide binding pocket is essential, as the presence of GDP or the absence of any nucleotide does not split the 55S mitoribosome (Tsuboi et al., 2009).

### Divergent mechanisms of fusidic acid (FA) resistance by EF-G1_mt_ and EF-G2_mt_

Fusidic Acid (FA) is a fusidane class antibiotic that is used to treat bacterial skin infections along with chronic bone and joint infections. It is effective against several species of gram-positive bacteria and is clinically used to treat methicillin-resistant *S. aureus* (MRSA). FA prevents the release of EF-G•GDP from the 70S ribosome after GTP hydrolysis by preventing the switch II from attaining its GDP-bound conformation (Gao et al., 2009). Prior structural studies have demonstrated that FA binds in an interdomain pocket between the G domain, domain II and domain III of EF-G (Gao et al., 2009; Zhou et al., 2013). Stable binding of FA requires the switch I region to be disordered (Gao et al., 2009) because an ordered switch I would overlap with the binding site of FA. Biochemical studies have shown that a substantially higher concentration (10-to 100-fold) of FA is needed to inhibit the activity of EF-G1_mt_ during mitochondrial elongation (Bhargava et al., 2004; Chung and Spremulli, 1990). Recent cryo-EM structures of EF-G1_mt_ bound to the 55S mitoribosomes have presented the switch I in a well-defined conformation (Koripella et al., 2020; Kummer and Ban, 2020) (Fig. 6A), in contrast to bacterial 70S•EF-G complexes where the density for switch I has been consistently poorly resolved (Fig. 6B) (Chen et al., 2013; Gao et al., 2009; Tourigny et al., 2013; Zhou et al., 2013, 2014). It was proposed that the increased resistance observed for EF-G1_mt_ towards FA resulted from a small insertion in the switch I region of EF-G1_mt_ (Kummer and Ban, 2020). The two positively charged lysine residues (Lys80 and Lys82) in this insertion form salt bridges with the negatively charged phosphate backbone of the SRL from the 39S subunit (Fig. 6B), and hence were hypothesized to confer additional stability to the switch I of EF-G1_mt_ (Kummer and Ban, 2020). Biochemical evidence showed EF-G1_mt_ to be more resistant towards FA than its bacterial counterpart (Bhargava et al., 2004; Chung and Spremulli, 1990).

**Fig. 6.**
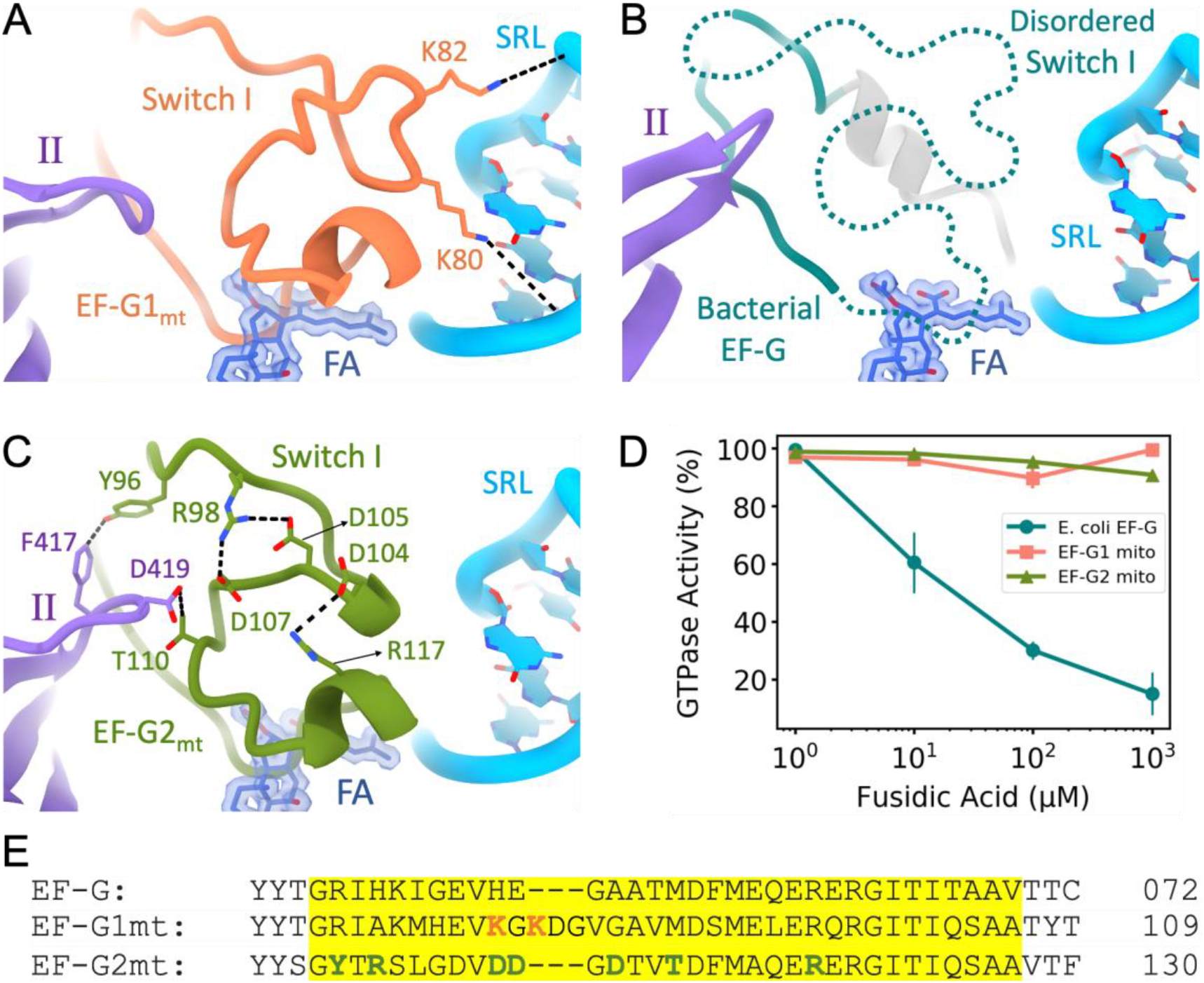
EF-G1_mt_ and EF-G2_mt_ follow diverse mechanisms to render resistance to the antibiotic fusidic acid (FA). **(A)** Stabilized Switch I region (salmon) (Koripella et al., 2020) in EF-G1_mt_ blocks FA (dark blue) from accessing its binding site. Switch I stability is achieved by the presence of two unique lysine residues (K80 and K82) that strongly interact with the phosphates of SRL by forming salt bridges (light blue) (Kummer and Ban, 2020). **(B)** The absence of these lysine residues in the Switch I region (dark cyan) of bacterial EF-G disorders the Switch I (Gao et al., 2009) and hence makes it susceptible to FA binding and inhibition. **(C)** In EF-G2_mt_, the primary aa sequence of switch I (green) is highly altered enabling the formation of three salt bridges within its switch I, thereby stabilizing it. Furthermore, strong interactions are observed between the switch I and domain II (purple) in EF-G2_mt_. **(D)** EF-G1_mt_ (salmon) and EF-G2_mt_ (green) show strong resistance even at high concentrations of FA, while the bacterial EF-G (dark cyan) is susceptible to FA inhibition even at low concentrations. **(E)** The aa sequence alignment of the switch I region (highlighted in yellow) in three EF-Gs. The lysine residues shown in salmon are unique to mammalian EF-G1_mt_ and confer stability to switch I by interacting with the SRL. Residues shown in green (except R117) are unique to EF-G2_mt_ and confer stability to switch I by forming salt bridges within the switch I region.

FA is known to inhibit both the translocation and the ribosome recycling steps in bacteria, though there is conflicting evidence on which step of translation is primarily targeted by FA (Borg et al., 2015; Borg et al., 2016; Savelsbergh et al., 2009). Comparison of the FA binding pocket between the bacterial EF-G, EF-G1_mt_ and EF-G2_mt_ revealed that key aa residues (Fig. S8A) reported to be necessary for the stable binding of FA (Gao et al., 2009; Zhou et al., 2013) are highly conserved (Fig. S8C), thereby suggesting a similar binding mechanism for FA for all the three EF-Gs. The three aa insertion in EF-G1_mt_ that confers resistance to FA is not present in EF-G2_mt_ and the corresponding region in EF-G2_mt_ does not contain any positively charged aa residues (Fig. 6E) that could strongly interact with the SRL. However, the cryo-EM map of EF-G2_mt_ shows a well-resolved density for the switch I region (Fig. S8B) and enabled its modelling (Fig. 6C), indicating an alternative mechanism for the stabilization of switch I in EF-G2_mt_.

Sequence alignment showed that the switch I region composition in EF-G2_mt_ is significantly different as compared to EF-G1_mt_ and the bacterial EF-G (Fig. 6E). Three new salt bridge interactions were identified within the switch I of EF-G2_mt_. Arg98 forms the first two salt bridges by pairing with Asp105 and Asp107 respectively, while Arg117 and Asp104 are involved in the formation of the third salt bridge (Fig. 6C). Furthermore, stronger interactions are observed between the switch I and domain II in EF-G2_mt_ compared to EF-G1_mt_. Thr110 from switch I is placed in the close proximity of Asp419 of domain II with the possibility of hydrogen bond formation (Fig. 6C) while the corresponding interaction in EF-G1_mt_ is between Asp404 and Val88, a much weaker interaction with no possibility of hydrogen bond formation. There is also potential for a tight T-stacking interaction between Phe417 and Tyr96 in EF-G2_mt_ (Fig. 6C), while the corresponding residues in EF-G1_mt_ being His402 and Arg72 offer no such interaction. Overall, through a combination of internal salt bridges and additional contacts with domain II, switch I gets highly stabilized in EF-G2_mt_. Since a stabilized switch I region occludes the binding site of FA (Fig. 6A,B), EF-G2_mt_ is expected to exhibit strong resistance towards FA in the lines of EF-G1_mt_. To test this hypothesis, we measured the GTPase activity of EF-G2_mt_ alongside EF-G1mt and *E. coli* EF-G in the presence of FA under multiple-turnover conditions (see Supplemental Materials and Methods). Our data shows that while 1μM FA has no effect on the GTPase activity in *E. coli*, significant inhibition is observed at higher concentrations of FA. In contrast, FA has almost no effect on the GTPase activity of either EF-G1_mt_ or EF-G2_mt_ even up to 10 mM FA. Our results are consistent with the earlier finding that EF-G1_mt_ is highly resistant to FA compared to the bacterial EF-G (Bhargava et al., 2004; Chung and Spremulli, 1990), and also consistent with recent finding that showed EF-G2_mt_ is not susceptible to inhibition by FA (Lee et al., 2020). Our results and analysis suggest FA resistance in EF-G2_mt_ occurs by a structural mechanism fundamentally different from that in EF-G1_mt_.

In conclusion, structures of three distinct functional states formed during the process of human mitoribosome recycling are presented (**Fig. 1**). A previous biochemical finding that GTP hydrolysis is not required for the RRF_mt_•EF-G2_mt_-mediated splitting of the post-termination mitoribosomal complex is corroborated. We also show that a mito-specific segment of the RRF_mt_’s NTE compensates for the slightly reduced size of the H69 within the 16S rRNA of the mitoribosomal large subunit (**Fig. 2**). This is the first evidence showing a translational factor compensating for an rRNA segment lost during the evolution of the mitoribosomal translation machinery. Our structures reveal how RRF_mt_’s domain II and EF-G2_mt_ domain IV directly help in disrupting the central inter-subunit bridge, B2a (**Fig. 3**), and suggest how the dynamics of the interactions among the uL11m, CTD of uL12m, and the G’domain of EF-G2_mt_ alters between elongation and recycling steps (**Fig. 4**). Structural analysis of domain IV of EF-G1_mt_ and EF-G2_mt_ explains their specific roles in two distinct steps of elongation and recycling, respectively (**Fig. 5**). Analysis of their GTPase domains complemented by GTPase assays reveal two distinct mechanisms of antibiotic fusidic acid resistance adopted by two homologous GTPases (**Fig. 6**). These observations help highlight the unique features of the main steps of human mitoribosomal recycling (**Fig. 7**). Future studies using time-resolved cryo-EM should help to resolve the short-lived intermediates that form during the transition from Class I to Class III states, which would help further characterize the functional roles of mito-specific segments of the two translational factors, RRF_mt_ and EF-G2_mt_.

**Fig. 7.**
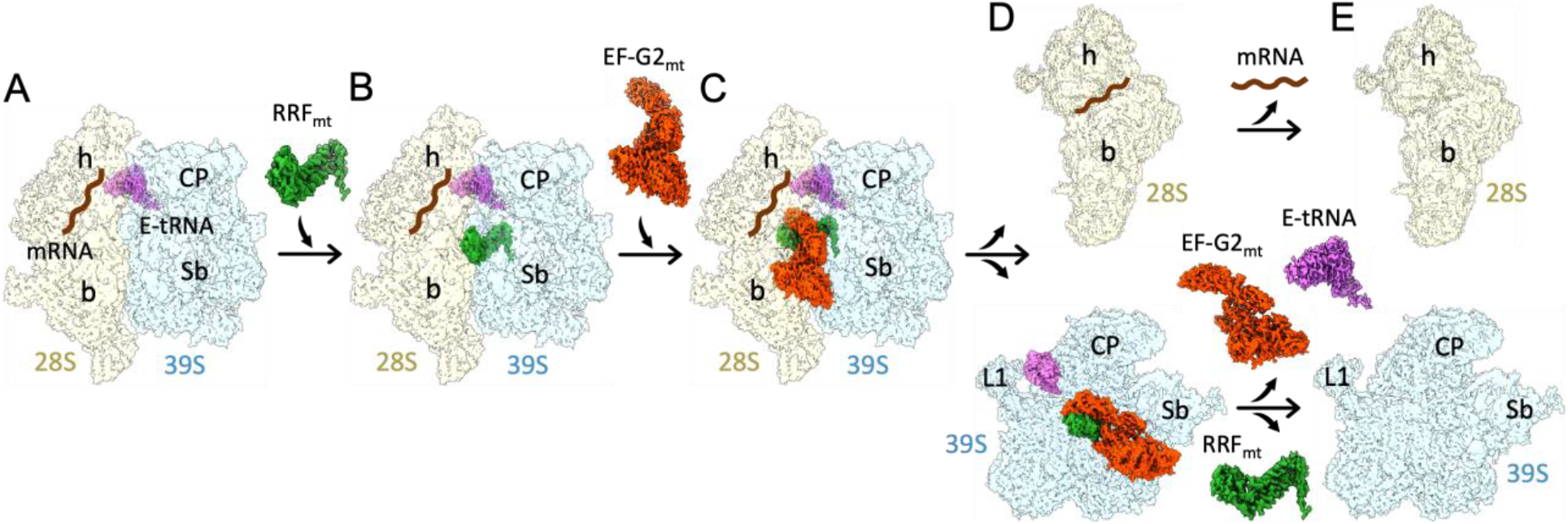
A sequence of main events in human mitochondrial ribosome recycling suggested by structures in this study. **(A)** The model post-termination mitoribosomal complex (PoTC), with mRNA and pe/E-state tRNAmt (Koripella et al., 2019). **(B)** Binding of RRF_mt_ locks the mitoribosome in a partially destabilized state with rotated 28S subunit (Fig. 1A, Fig. S…). The mito-specific N-terminus of the RRF_mt_’s NTE occupies a unique site near the bridge B2a region (Fig. 2). **(C)** Subsequent binding of EF-G2_mt_ further destabilizes the 55S complex with disruption of additional inter-subunit bridges, in a fast reaction leading to multiple short-lived intermediate states, with the 28S subunit present in multiple orientations relative to the 39S subunit (Fig. 1B). **(D)** 28S and 39S subunits are dissociated. While the 28S subunit still carries mRNA, the 39S subunit carries RRF_mt_, EF-G2_mt_, and surprisingly a tRNAmt in the 39S subunit’s E-site region (Fig. 1C). This is in sharp contrast to the tRNA dissociation mechanism in eubacterial ribosome recycling, where tRNA goes with the small 30S ribosomal subunit. **(E)** Depiction of final recycling products, steps of ligand release between panels D and E are yet to be characterized.

## Acknowledgements

We thank Linda Spremulli (UNC), and Robert Lightowlers and Zofia Chrzanowska-Lightowlers (New Castle, UK) for sharing their RRF_mt_ and EF-G2_mt_ clones. We thank Manjuli Sharma for helpful suggestions and comments during the course of this study and on the manuscript. We acknowledge the use of the Wadsworth Center’s Media and Culture core facility, for help in producing large volumes of HEK293S GnTI cells, and the Wadsworth Center’s and New York Structural Biology Center’s (NYSBC’s) EM facilities. NYSBC EM facilities are supported by grants from the Simons Foundation (349247), NYSTAR, the NIH (GM103310) and the Agouron Institute (F00316). This work was supported by the NIH grant R01 GM61576 (to R.K.A.).

## Author Contributions

RKA conceived this study. PK purified human mitoribosomes, human RRF_mt_ and EF-G2_mt_, and RKK prepared the human 55S PoTC•RRF_mt_•EF-G2_mt_ complex. RKK and NKB collected cryo-EM data, and RKK and RKA performed image processing. RKK performed molecular modeling, using assistance from AD, EKA and NKB. AD, EKA and PK performed the biochemical assays. RKK, AD, and RKA analyzed the data and wrote the manuscript. All authors read and approved the manuscript. Authors declare no competing financial interest in this work.

## Data Deposition

The cryo-EM maps and atomic coordinates described in this study have been deposited in the Electron Microscopy and PDB Data Bank (wwPDB.org) under accession codes EMD-23096 and PDB ID: 7L08 for the RRF_mt_-bound human 55S mitoribosome (Complex I), EMD-23114 for the RRF_mt_•EF-G2_mt_-bound human 55S mitoribosome (Complex II), and EMD-23121 and PDB ID: 7L20 for the RRF_mt_•EF-G2_mt_-bound human 39S mitoribosome (Complex III).

## References

AEvarsson, A., Brazhnikov, E., Garber, M., Zheltonosova, J., Chirgadze, Y., al-Karadaghi, S., Svensson, L.A., and Liljas, A. (1994). Three-dimensional structure of the ribosomal translocase: elongation factor G from Thermus thermophilus. EMBO J 13, 3669–3677.

Agrawal, R.K., Sharma, M.R., Kiel, M.C., Hirokawa, G., Booth, T.M., Spahn, C.M., Grassucci, R.A., Kaji, A., and Frank, J. (2004). Visualization of ribosome-recycling factor on the Escherichia coli 70S ribosome: functional implications. Proc Natl Acad Sci U S A 101, 8900–8905.

Aibara, S., Singh, V., Modelska, A., and Amunts, A. (2020). Structural basis of mitochondrial translation. Elife 9.

Amunts, A., Brown, A., Toots, J., Scheres, S.H.W., and Ramakrishnan, V. (2015). Ribosome. The structure of the human mitochondrial ribosome. Science 348, 95–98.

Barat, C., Datta, P.P., Raj, V.S., Sharma, M.R., Kaji, H., Kaji, A., and Agrawal, R.K. (2007). Progression of the ribosome recycling factor through the ribosome dissociates the two ribosomal subunits. Mol Cell 27, 250–261.

Bhargava, K., Templeton, P., and Spremulli, L.L. (2004). Expression and characterization of isoform 1 of human mitochondrial elongation factor G. Protein Expr Purif 37, 368–376.

Boerema, A.P., Aibara, S., Paul, B., Tobiasson, V., Kimanius, D., Forsberg, B.O., Wallden, K., Lindahl, E., and Amunts, A. (2018). Structure of the chloroplast ribosome with chl-RRF and hibernation-promoting factor. Nat Plants 4, 212–217.

Borg, A., Holm, M., Shiroyama, I., Hauryliuk, V., Pavlov, M., Sanyal, S., and Ehrenberg, M. (2015). Fusidic acid targets elongation factor G in several stages of translocation on the bacterial ribosome. J Biol Chem 290, 3440–3454.

Borg, A., Pavlov, M., and Ehrenberg, M. (2016). Complete kinetic mechanism for recycling of the bacterial ribosome. RNA 22, 10–21.

Brown, A., Amunts, A., Bai, X.C., Sugimoto, Y., Edwards, P.C., Murshudov, G., Scheres, S.H.W., and Ramakrishnan, V. (2014). Structure of the large ribosomal subunit from human mitochondria. Science 346, 718–722.

Budkevich, T.V., Giesebrecht, J., Behrmann, E., Loerke, J., Ramrath, D.J., Mielke, T., Ismer, J., Hildebrand, P.W., Tung, C.S., Nierhaus, K.H., et al. (2014). Regulation of the mammalian elongation cycle by subunit rolling: a eukaryotic-specific ribosome rearrangement. Cell 158, 121–131.

Chen, Y., Feng, S., Kumar, V., Ero, R., and Gao, Y.G. (2013). Structure of EF-G-ribosome complex in a pretranslocation state. Nat Struct Mol Biol 20, 1077–1084.

Chen, Y., Koripella, R.K., Sanyal, S., and Selmer, M. (2010). Staphylococcus aureus elongation factor G--structure and analysis of a target for fusidic acid. FEBS J 277, 3789–3803.

Christian, B.E., and Spremulli, L.L. (2012). Mechanism of protein biosynthesis in mammalian mitochondria. Biochim Biophys Acta 1819, 1035–1054.

Chung, H.K., and Spremulli, L.L. (1990). Purification and characterization of elongation factor G from bovine liver mitochondria. J Biol Chem 265, 21000–21004.

Connell, S.R., Takemoto, C., Wilson, D.N., Wang, H., Murayama, K., Terada, T., Shirouzu, M., Rost, M., Schuler, M., Giesebrecht, J., et al. (2007). Structural basis for interaction of the ribosome with the switch regions of GTP-bound elongation factors. Mol Cell 25, 751–764.

Czworkowski, J., Wang, J., Steitz, T.A., and Moore, P.B. (1994). The crystal structure of elongation factor G complexed with GDP, at 2.7 A resolution. EMBO J 13, 3661–3668.

Datta, P.P., Sharma, M.R., Qi, L., Frank, J., and Agrawal, R.K. (2005). Interaction of the G’ domain of elongation factor G and the C-terminal domain of ribosomal protein L7/L12 during translocation as revealed by cryo-EM. Mol Cell 20, 723–731.

Desai, N., Brown, A., Amunts, A., and Ramakrishnan, V. (2017). The structure of the yeast mitochondrial ribosome. Science 355, 528–531.

Diaconu, M., Kothe, U., Schlunzen, F., Fischer, N., Harms, J.M., Tonevitsky, A.G., Stark, H., Rodnina, M.V., and Wahl, M.C. (2005). Structural basis for the function of the ribosomal L7/12 stalk in factor binding and GTPase activation. Cell 121, 991–1004.

Dunkle, J.A., Wang, L., Feldman, M.B., Pulk, A., Chen, V.B., Kapral, G.J., Noeske, J., Richardson, J.S., Blanchard, S.C., and Cate, J.H. (2011). Structures of the bacterial ribosome in classical and hybrid states of tRNA binding. Science 332, 981–984.

Eberly, S.L., Locklear, V., and Spremulli, L.L. (1985). Bovine mitochondrial ribosomes. Elongation factor specificity. J Biol Chem 260, 8721–8725.

Frank, J., and Agrawal, R.K. (2000). A ratchet-like inter-subunit reorganization of the ribosome during translocation. Nature 406, 318–322.

Frank, J., and Agrawal, R.K. (2001). Ratchet-like movements between the two ribosomal subunits: their implications in elongation factor recognition and tRNA translocation. Cold Spring Harb Symp Quant Biol 66, 67–75.

Freistroffer, D.V., Pavlov, M.Y., MacDougall, J., Buckingham, R.H., and Ehrenberg, M. (1997). Release factor RF3 in E.coli accelerates the dissociation of release factors RF1 and RF2 from the ribosome in a GTP-dependent manner. EMBO J 16, 4126–4133.

Fu, Z., Kaledhonkar, S., Borg, A., Sun, M., Chen, B., Grassucci, R.A., Ehrenberg, M., and Frank, J. (2016). Key Intermediates in Ribosome Recycling Visualized by Time-Resolved Cryoelectron Microscopy. Structure 24, 2092–2101.

Fujiwara, T., Ito, K., and Nakamura, Y. (2001). Functional mapping of ribosome-contact sites in the ribosome recycling factor: a structural view from a tRNA mimic. RNA 7, 64–70.

Fujiwara, T., Ito, K., Nakayashiki, T., and Nakamura, Y. (1999). Amber mutations in ribosome recycling factors of Escherichia coli and Thermus thermophilus: evidence for C-terminal modulator element. FEBS Lett 447, 297–302.

Gao, N., Zavialov, A.V., Ehrenberg, M., and Frank, J. (2007). Specific interaction between EF-G and RRF and its implication for GTP-dependent ribosome splitting into subunits. J Mol Biol 374, 1345–1358.

Gao, N., Zavialov, A.V., Li, W., Sengupta, J., Valle, M., Gursky, R.P., Ehrenberg, M., and Frank, J. (2005). Mechanism for the disassembly of the posttermination complex inferred from cryo-EM studies. Mol Cell 18, 663–674.

Gao, Y.G., Selmer, M., Dunham, C.M., Weixlbaumer, A., Kelley, A.C., and Ramakrishnan, V. (2009). The structure of the ribosome with elongation factor G trapped in the posttranslocational state. Science 326, 694–699.

Gray, M.W., Burger, G., and Lang, B.F. (1999). Mitochondrial evolution. Science 283, 1476–1481.

Greber, B.J., Bieri, P., Leibundgut, M., Leitner, A., Aebersold, R., Boehringer, D., and Ban, N. (2015). Ribosome. The complete structure of the 55S mammalian mitochondrial ribosome. Science 348, 303–308.

Hammarsund, M., Wilson, W., Corcoran, M., Merup, M., Einhorn, S., Grander, D., and Sangfelt, O. (2001). Identification and characterization of two novel human mitochondrial elongation factor genes, hEFG2 and hEFG1, phylogenetically conserved through evolution. Hum Genet 109, 542–550.

Helgstrand, M., Mandava, C.S., Mulder, F.A., Liljas, A., Sanyal, S., and Akke, M. (2007). The ribosomal stalk binds to translation factors IF2, EF-Tu, EF-G and RF3 via a conserved region of the L12 C-terminal domain. J Mol Biol 365, 468–479.

Hirokawa, G., Kiel, M.C., Muto, A., Selmer, M., Raj, V.S., Liljas, A., Igarashi, K., Kaji, H., and Kaji, A. (2002). Post-termination complex disassembly by ribosome recycling factor, a functional tRNA mimic. EMBO J 21, 2272–2281.

Hirokawa, G., Nijman, R.M., Raj, V.S., Kaji, H., Igarashi, K., and Kaji, A. (2005). The role of ribosome recycling factor in dissociation of 70S ribosomes into subunits. RNA 11, 1317–1328.

Imai, H., Uchiumi, T., and Kodera, N. (2020). Direct visualization of translational GTPase factor pool formed around the archaeal ribosomal P-stalk by high-speed AFM. Proc Natl Acad Sci U S A.

Ito, K., Ebihara, K., Uno, M., and Nakamura, Y. (1996). Conserved motifs in prokaryotic and eukaryotic polypeptide release factors: tRNA-protein mimicry hypothesis. Proc Natl Acad Sci U S A 93, 5443–5448.

Ito, K., Fujiwara, T., Toyoda, T., and Nakamura, Y. (2002). Elongation factor G participates in ribosome disassembly by interacting with ribosome recycling factor at their tRNA-mimicry domains. Mol Cell 9, 1263–1272.

Iwakura, N., Yokoyama, T., Quaglia, F., Mitsuoka, K., Mio, K., Shigematsu, H., Shirouzu, M., Kaji, A., and Kaji, H. (2017). Chemical and structural characterization of a model Post-Termination Complex (PoTC) for the ribosome recycling reaction: Evidence for the release of the mRNA by RRF and EF-G. PLoS One 12, e0177972.

Janosi, L., Mottagui-Tabar, S., Isaksson, L.A., Sekine, Y., Ohtsubo, E., Zhang, S., Goon, S., Nelken, S., Shuda, M., and Kaji, A. (1998). Evidence for in vivo ribosome recycling, the fourth step in protein biosynthesis. EMBO J 17, 1141–1151.

Kaji, A., Kiel, M.C., Hirokawa, G., Muto, A.R., Inokuchi, Y., and Kaji, H. (2001). The fourth step of protein synthesis: disassembly of the posttermination complex is catalyzed by elongation factor G and ribosome recycling factor, a near-perfect mimic of tRNA. Cold Spring Harb Symp Quant Biol 66, 515–529.

Karimi, R., Pavlov, M.Y., Buckingham, R.H., and Ehrenberg, M. (1999). Novel roles for classical factors at the interface between translation termination and initiation. Mol Cell 3, 601–609.

Kaushal, P.S., Sharma, M.R., Booth, T.M., Haque, E.M., Tung, C.S., Sanbonmatsu, K.Y., Spremulli, L.L., and Agrawal, R.K. (2014). Cryo-EM structure of the small subunit of the mammalian mitochondrial ribosome. Proc Natl Acad Sci U S A 111, 7284–7289.

Kim, K.K., Min, K., and Suh, S.W. (2000). Crystal structure of the ribosome recycling factor from Escherichia coli. EMBO J 19, 2362–2370.

Kisselev, L., Ehrenberg, M., and Frolova, L. (2003). Termination of translation: interplay of mRNA, rRNAs and release factors? EMBO J 22, 175–182.

Kolesnikov, A.V., and Gudkov, A.T. (2003). [Mutational analysis of the functional role of the loop region in the elongation factor G fourth domain in the ribosomal translocation]. Mol Biol (Mosk) 37, 719–725.

Koripella, R.K., Sharma, M.R., Bhargava, K., Datta, P.P., Kaushal, P.S., Keshavan, P., Spremulli, L.L., Banavali, N.K., and Agrawal, R.K. (2020). Structures of the human mitochondrial ribosome bound to EF-G1 reveal distinct features of mitochondrial translation elongation. Nat Commun 11, 3830.

Koripella, R.K., Sharma, M.R., Haque, M.E., Risteff, P., Spremulli, L.L., and Agrawal, R.K. (2019a). Structure of Human Mitochondrial Translation Initiation Factor 3 Bound to the Small Ribosomal Subunit. iScience 12, 76–86.

Koripella, R.K., Sharma, M.R., Risteff, P., Keshavan, P., and Agrawal, R.K. (2019b). Structural insights into unique features of the human mitochondrial ribosome recycling. Proc Natl Acad Sci U S A 116, 8283–8288.

Kummer, E., and Ban, N. (2020). Structural insights into mammalian mitochondrial translation elongation catalyzed by mtEFG1. EMBO J 39, e104820.

Kummer, E., Leibundgut, M., Rackham, O., Lee, R.G., Boehringer, D., Filipovska, A., and Ban, N. (2018). Unique features of mammalian mitochondrial translation initiation revealed by cryo-EM. Nature 560, 263–267.

Lee, M., Matsunaga, N., Akabane, S., Yasuda, I., Ueda, T., and Takeuchi-Tomita, N. (2020). Reconstitution of mammalian mitochondrial translation system capable of correct initiation and long polypeptide synthesis from leaderless mRNA. Nucleic Acids Res.

Margus, T., Remm, M., and Tenson, T. (2007). Phylogenetic distribution of translational GTPases in bacteria. BMC Genomics 8, 15.

Nakano, H., Yoshida, T., Uchiyama, S., Kawachi, M., Matsuo, H., Kato, T., Ohshima, A., Yamaichi, Y., Honda, T., Kato, H., et al. (2003). Structure and binding mode of a ribosome recycling factor (RRF) from mesophilic bacterium. J Biol Chem 278, 3427–3436.

Pel, H.J., and Grivell, L.A. (1994). Protein synthesis in mitochondria. Mol Biol Rep 19, 183–194.

Peske, F., Rodnina, M.V., and Wintermeyer, W. (2005). Sequence of steps in ribosome recycling as defined by kinetic analysis. Mol Cell 18, 403–412.

Poole, E., and Tate, W. (2000). Release factors and their role as decoding proteins: specificity and fidelity for termination of protein synthesis. Biochim Biophys Acta 1493, 1–11.

Pulk, A., and Cate, J.H. (2013). Control of ribosomal subunit rotation by elongation factor G. Science 340, 1235970.

Rao, A.R., and Varshney, U. (2001). Specific interaction between the ribosome recycling factor and the elongation factor G from Mycobacterium tuberculosis mediates peptidyl-tRNA release and ribosome recycling in Escherichia coli. EMBO J 20, 2977–2986.

Ratje, A.H., Loerke, J., Mikolajka, A., Brunner, M., Hildebrand, P.W., Starosta, A.L., Donhofer, A., Connell, S.R., Fucini, P., Mielke, T., et al. (2010). Head swivel on the ribosome facilitates translocation by means of intra-subunit tRNA hybrid sites. Nature 468, 713–716.

Rorbach, J., Richter, R., Wessels, H.J., Wydro, M., Pekalski, M., Farhoud, M., Kuhl, I., Gaisne, M., Bonnefoy, N., Smeitink, J.A., et al. (2008). The human mitochondrial ribosome recycling factor is essential for cell viability. Nucleic Acids Res 36, 5787–5799.

Saikrishnan, K., Kalapala, S.K., Varshney, U., and Vijayan, M. (2005). X-ray structural studies of Mycobacterium tuberculosis RRF and a comparative study of RRFs of known structure. Molecular plasticity and biological implications. J Mol Biol 345, 29–38.

Savelsbergh, A., Rodnina, M.V., and Wintermeyer, W. (2009). Distinct functions of elongation factor G in ribosome recycling and translocation. RNA 15, 772–780.

Schuwirth, B.S., Borovinskaya, M.A., Hau, C.W., Zhang, W., Vila-Sanjurjo, A., Holton, J.M., and Cate, J.H. (2005). Structures of the bacterial ribosome at 3.5 A resolution. Science 310, 827–834.

Selmer, M., Al-Karadaghi, S., Hirokawa, G., Kaji, A., and Liljas, A. (1999). Crystal structure of Thermotoga maritima ribosome recycling factor: a tRNA mimic. Science 286, 2349–2352.

Seshadri, A., Samhita, L., Gaur, R., Malshetty, V., and Varshney, U. (2009). Analysis of the fusA2 locus encoding EFG2 in Mycobacterium smegmatis. Tuberculosis (Edinb) 89, 453–464.

Sharma, M.R., Kaushal, P.S., Gupta, M., Banavali, N.K., and Agrawal, R.K. (2013). Insights into Structural Basis of Mammalian Mitochondrial Translation. In Translation in Mitochondria and Other Organelles, D. A-M, ed. (Berlin: Springer), pp. 1–28.

Sharma, M.R., Koc, E.C., Datta, P.P., Booth, T.M., Spremulli, L.L., and Agrawal, R.K. (2003). Structure of the mammalian mitochondrial ribosome reveals an expanded functional role for its component proteins. Cell 115, 97–108.

Sharma, M.R., Wilson, D.N., Datta, P.P., Barat, C., Schluenzen, F., Fucini, P., and Agrawal, R.K. (2007). Cryo-EM study of the spinach chloroplast ribosome reveals the structural and functional roles of plastid-specific ribosomal proteins. Proc Natl Acad Sci U S A 104, 19315–19320.

Suematsu, T., Yokobori, S., Morita, H., Yoshinari, S., Ueda, T., Kita, K., Takeuchi, N., and Watanabe, Y. (2010). A bacterial elongation factor G homologue exclusively functions in ribosome recycling in the spirochaete Borrelia burgdorferi. Mol Microbiol 75, 1445–1454.

Tourigny, D.S., Fernandez, I.S., Kelley, A.C., and Ramakrishnan, V. (2013). Elongation factor G bound to the ribosome in an intermediate state of translocation. Science 340, 1235490.

Toyoda, T., Tin, O.F., Ito, K., Fujiwara, T., Kumasaka, T., Yamamoto, M., Garber, M.B., and Nakamura, Y. (2000). Crystal structure combined with genetic analysis of the Thermus thermophilus ribosome recycling factor shows that a flexible hinge may act as a functional switch. RNA 6, 1432–1444.

Tsuboi, M., Morita, H., Nozaki, Y., Akama, K., Ueda, T., Ito, K., Nierhaus, K.H., and Takeuchi, N. (2009). EF-G2_mt_ is an exclusive recycling factor in mammalian mitochondrial protein synthesis. Mol Cell 35, 502–510.

Weixlbaumer, A., Petry, S., Dunham, C.M., Selmer, M., Kelley, A.C., and Ramakrishnan, V. (2007). Crystal structure of the ribosome recycling factor bound to the ribosome. Nat Struct Mol Biol 14, 733–737.

Yassin, A.S., Haque, M.E., Datta, P.P., Elmore, K., Banavali, N.K., Spremulli, L.L., and Agrawal, R.K. (2011). Insertion domain within mammalian mitochondrial translation initiation factor 2 serves the role of eubacterial initiation factor 1. Proc Natl Acad Sci U S A 108, 3918–3923.

Yokoyama, T., Shaikh, T.R., Iwakura, N., Kaji, H., Kaji, A., and Agrawal, R.K. (2012). Structural insights into initial and intermediate steps of the ribosome-recycling process. EMBO J 31, 1836–1846.

Yoshida, T., Uchiyama, S., Nakano, H., Kashimori, H., Kijima, H., Ohshima, T., Saihara, Y., Ishino, T., Shimahara, H., Yoshida, T., et al. (2001). Solution structure of the ribosome recycling factor from Aquifex aeolicus. Biochemistry 40, 2387–2396.

Zavialov, A.V., Hauryliuk, V.V., and Ehrenberg, M. (2005). Splitting of the posttermination ribosome into subunits by the concerted action of RRF and EF-G. Mol Cell 18, 675–686.

Zavialov, A.V., Mora, L., Buckingham, R.H., and Ehrenberg, M. (2002). Release of peptide promoted by the GGQ motif of class 1 release factors regulates the GTPase activity of RF3. Mol Cell 10, 789–798.

Zhang, D., Yan, K., Zhang, Y., Liu, G., Cao, X., Song, G., Xie, Q., Gao, N., and Qin, Y. (2015). New insights into the enzymatic role of EF-G in ribosome recycling. Nucleic Acids Res 43, 10525–10533.

Zhang, Y., and Spremulli, L.L. (1998). Identification and cloning of human mitochondrial translational release factor 1 and the ribosome recycling factor. Biochim Biophys Acta 1443, 245–250.

Zhou, D., Tanzawa, T., Lin, J., and Gagnon, M.G. (2020). Structural basis for ribosome recycling by RRF and tRNA. Nat Struct Mol Biol 27, 25–32.

Zhou, J., Lancaster, L., Donohue, J.P., and Noller, H.F. (2013). Crystal structures of EF-G-ribosome complexes trapped in intermediate states of translocation. Science 340, 1236086.

Zhou, J., Lancaster, L., Donohue, J.P., and Noller, H.F. (2014). How the ribosome hands the A-site tRNA to the P site during EF-G-catalyzed translocation. Science 345, 1188–1191.

