## Supplementary material for "Structures of the human mitochondrial ribosome recycling complexes reveal distinct mechanisms of recycling and antibiotic resistance": Materials Methods, Figures, and Table

### Materials and Methods:

#### Purification of 55S mitoribosomes and $RRF_{mt}$:

The 55S mitoribosomes were purified from human embryonic kidney cells lacking N-acetyl-glucosaminyltransferase I (HEK293S GnTI) according to protocols described previously (Koripella et al., 2019). SUMO-tagged human $RRF_{mt}$ that was cloned into pET24(C+) plasmid was overexpressed in Rosetta2 cell lines and then purified using affinity chromatography as described earlier (Koripella et al., 2019).

#### Overexpression and purification of EF-$G2_{mt}$:

The GST-tagged EF-$G2_{mt}$ was cloned into pGEX6.1 vector and over-expressed in Rossetta (pLysS+RARE) cell lines. Cells were grown in LB media with 100 ug/ml ampicillin until 0.6 O.D. and protein over-expression was induced by adding 100 mM isopropyl-1-thio-D-galactopyranoside (IPTG). The cell culture was left overnight at 16 °C for optimal protein yields. The cells were pelleted in a JLA 10.5 rotor at 5000 rpm for 30 minutes and were shock-frozen in liquid nitrogen and stored at -80 °C. The frozen cells were resuspended in lysis buffer (1XPBS-pH7.5, 10 mM MgCl2, 1 mM phenylmethyl sulphonyl fluoride (PMSF) and 1 mM DTT). After sonication and DNase I (5 μg/mL) treatment, the lysate were centrifuged at 14,000 rpm for 30 minutes to remove cell debris. The supernatant was passed through GSTrap™ HP column equilibrated with binding buffer 1XPBS-pH7.5 and 1 mM DTT. The GST-tag was cleaved from EF-$G2_{mt}$ by loading 3C pression protease (gifted by Dr. Hongmin Li's lab, Wadsworth center, USA) along with cleavage buffer (50 mM Tris-HCl-pH 7.5, 150 mM NaCl, and 1 mM DTT). After incubating the EF-$G2_{mt}$ with 3C precision protease overnight, relatively pure EF-$G2_{mt}$ was eluted by passing the elution buffer (50 mM Tris-HCl-pH 7.5, 150 mM NaCl and 1mM DTT). To obtain high-level purity, EF-$G2_{mt}$ was further passed through an anion exchange HiTrap Q HP column (GE healthcare, USA) and pure protein was eluted by running a gradient with 20 mM Tris-HCl-pH 7.5 and 500 mM NaCl.

#### Preparation of the 55S•$RRF_{mt}$•EF-$G2_{mt}$•GMPPCP complex:

50 μM puromycin and 150 nM 55S mitoribosomes were mixed in HEPES polymix buffer (5 mM HEPES-KOH pH 7.5, 100 mM KCl, 20 mM MgOAc, 5 mM $NH_4Cl$, 0.5 mM $CaCl_2$, 1 mM DTT, 1 mM spermidine, and 8 mM putrescine) and incubated for 10 minutes at 37 °C to obtain the

model post-termination (PoTC) complex. 15 µM $RRF_{mt}$ was added to this reaction mixture of PoTC and incubated for an additional 5 minutes at 37°C to obtain the 55S•$RRF_{mt}$ complex as described earlier (Koripella et al., 2019). 5 µM EF-$G2_{mt,}$ together with 200 µM GMPPCP, was added to the 55S•$RRF_{mt}$ complex and incubated for various timepoints (5 sec, 30 sec and 2 minutes) at 37°C to obtain the 55S•$RRF_{mt}$•EF-$G2_{mt}$•GMPPCP complex, which was immediately utilized for the cryo-EM grid preparation.

**Cryo-electron microscopy and image processing:**

Home-made carbon was coated as a continuous layer (~50 Å thick) onto Quantifoil holey copper 1.2/1.3 grids, which were then glow-discharged for 30 seconds on a plasma sterilizer. 4 µl of the sample was loaded to each of the multiple grids, incubated for 15 seconds at 4 °C and 100 % humidity, and then blotted for 4 seconds before flash-freezing into the liquid ethane using a Vitrobot IV (FEI). Data was acquired on a Titan Krios electron microscope equipped with a Gatan K2 summit direct-electron detecting camera at 300 KV. A defocus range of -1.0 to -3.0 µm was used at a calibrated magnification of 22,500 X, yielding a pixel size of 1.0732 Å on the object scale. A dose rate of 8.25 electrons per pixel per second and an exposure time of 10 seconds resulted in a total dose of 71.6 $eÅ^{-2}$. CryoSPARC (Punjani et al., 2017) was employed for all the subsequent downstream data processing steps. After applying full-frame motion correction to all 50 movie frames corresponding to each of the 21,752 micrographs that were collected, 142 bad micrographs were deselected after determining their contrast transfer function (CTF) using CTFFIND4 (Rohou and Grigorieff, 2015). From the remaining 21,610 micrographs, a total of 5,046,906 particles were auto-picked which were then subjected to local motion correction and then 4,049,952 particles were retained. This step was followed by reference-free 2D classification which allowed us to further separate the good particles (1,140,751) from the bad particles (2,909,201) based on the 2D averages. Reference-based 3D classification was employed first to separate the particles into intact 55S mitoribosomes (162,354 particles), 39S subunits (419,699 particles), 28S subunits (235,896 particles) and poorly aligned particles (322,802 particles). To obtain more homogenous sub-populations, particles corresponding to the 55S mitoribosomes, 39S and 28S subunits were each subjected to additional rounds of 3D classification that finally yielded three stable classes representing three distinct functional states formed during the human mitoribosome recycling process. Two 55S mitoribosome classes, Class

I (93,212 particles) and Class II (28,929 particles), were finally refined to 3.49 and 3.91 Å, respectively, while the 39S subunit Class III (132,008 particles) was refined to 3.15 Å.

**Model building and optimization:**

Coordinates corresponding to the small and large subunits from our published human mitoribosome structures (Koripella et al., 2020) (PDB ID: 6VLZ) were docked independently as rigid bodies into the corresponding cryo-EM density maps of the Class I, Class II and Class III complexes using Chimera 1.14 (Pettersen EF 2004). To obtain optimal fitting into our cryo-EM densities, the models were subsequently refined in PHENIX (Adams PD 2010) using the "real-space refinement" function. Coordinates belonging to the human $RRF_{mt}$ (Koripella et al., 2019) were placed independently into the corresponding cryo-EM densities of all the three maps as rigid bodies using Chimera 1.14 (Pettersen et al., 2004) and the models were further real-space refined in PHENIX (Adams et al., 2010) to achieve optimal accommodation into the cryo-EM densities. The primary aa sequence of EF-$G2_{mt}$ was submitted to the I-TASSER server (Roy et al., 2010) to generate the initial EF-$G2_{mt}$ homology model that was used to interpret the corresponding cryo-EM density. Segments in the homology model that do not fully accommodate into the corresponding EF-$G2_{mt}$ density were modeled *de novo* using Chimera 1.14 (Pettersen et al., 2004) and COOT (Emsley et al., 2010). For the final optimization of the model into the cryo-EM density, the "Real-space refinement" function in PHENIX (Adams et al., 2010) was utilized. Validation reports for all the models were obtained from the Molprobity server (Chen et al., 2010) and the overall statistics of EM reconstruction and molecular modelling are listed in Table S1.

**GTPase-Glo assay:**

GTPase activity was measured using the GTPase-Glo™ Assay by Promega and carried out as described (Mondal et al., 2015). In brief, a 10 µl reaction consisting of 0.1 µM ribosomes, 0.25 µM either bacterial (*E. coli*) EF-G or EF-$G1_{mt}$ or EF-$G2_{mt}$ and 1 µM GTP were incubated at room temperature in GTPase-Glo™ buffer in the absence and presence of varying amounts of fusidic acid (FA) for 1 hour. 10 µl of Reconstituted GTPase-Glo™ reagent was added to the reaction and left shaking at room temperature for 30 minutes. Finally, 20 µL of detection reagent was added, incubated for 10 minutes at room temperature and luminescence was measured in a

Turner Biosystems Veritas™ Microplate Luminometer after 10 minutes. A negative control that contained only 1 µM GTP and a positive control that contained no FA were used each time. Luminescence was measured at FA concentrations of 1 µM, 10 µM, 100 µM, and 1000 µM. Data was collected in duplicates and each duplicate dataset was normalized between 0 and 1 using the equation $x_{norm} = \frac{x - x_{min}}{x_{max} - x_{min}}$. As GTPase activity and luminescence measured had an inverse relationship, % GTPase activity was calculated with the formula $\%\ GTPase\ Activity = (1 - x_{norm}) * 100$. The values at each point were then averaged and standard deviation was calculated. The plot was generated using Python library Matplotlib.

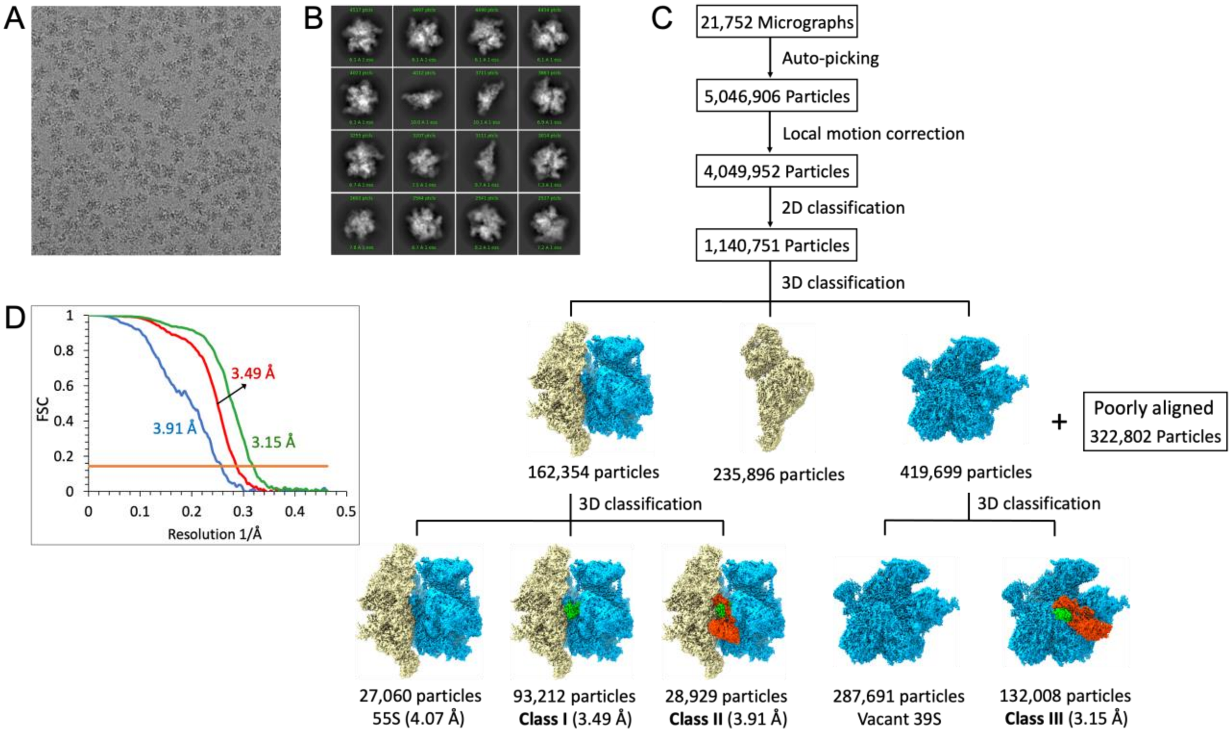


**Fig. S1. Image processing of the human mitochondrial ribosome recycling complexes.** (**A**) A typical electron micrograph obtained for the human mitoribosomal 55S•RRF$_{mt}$•EF-G2$_{mt}$•GMPPCP complex. (**B**) Representative two-dimensional (2D) class averages used in initial three-dimensional (3D) reconstructions. (**C**) Flow-chart showing results of 3D classifications and refinements. The selected 2D averages (1,140,751 particles) were subjected to several rounds of reference-based 3D classification to separate the intact 55S mitoribosomes (162,354 particles) from the 28S subunits (235,896 particles), the 39S subunits (419,699 particles) and the poorly aligned images (322,802 particles). Further classification of the 55S mitoribosomes yielded three stable classes. The 55S maps that contained bound ligands were refined to 3.49 Å (Class I) and 3.91 Å (Class II). Particles corresponding to the 39S subunit were subjected to additional rounds of 3D classification that finally yielded a stable 39S class (Class III) bound with both factors, RRF$_{mt}$ and EF-G2$_{mt}$. This 39S class was refined to 3.15 Å. **(D)** Fourier-shell correlation (FSC) plots of the Classs I (red), Class II (blue) and Class III (green) complexes.

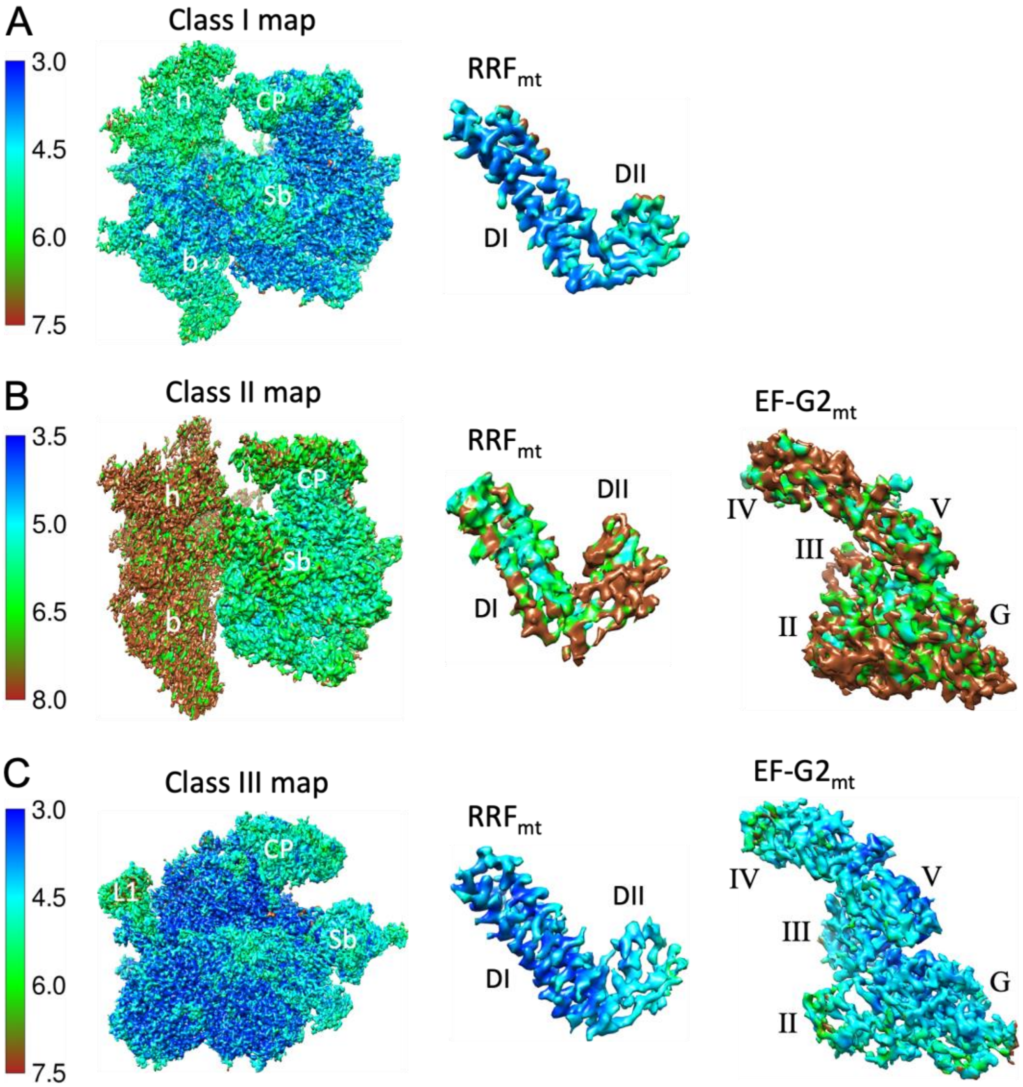


**Fig. S2. Local surface resolution of the human mitoribosome recycling complexes. (A-C)** The Left panels show the local resolution of Class I, Class II and Class III maps, respectively. The middle panels and the right panels show the local resolution of $RRF_{mt}$ and $EF\text{-}G_{mt}$ components extracted from the corresponding cryo-EM maps displayed in the left panels.

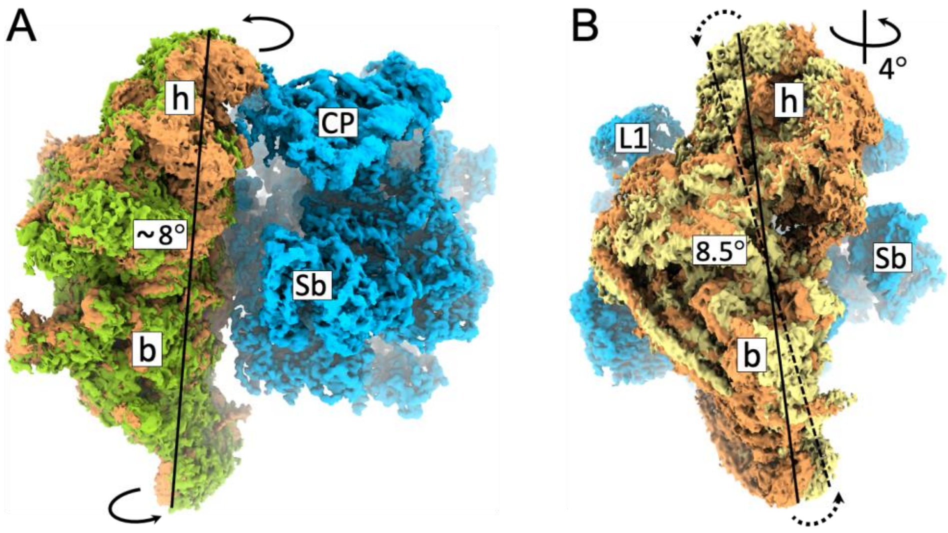


**Fig. S3. Conformational states of the 28S subunit in the Class I and Class II mitoribosomal recycling complexes.** (**A**) Comparison of our factor-free 55S cryo-EM map with the published factor-free human 55S mitoribosome (Amunts et al., 2015) (light brown) showed that 28S subunit (light green) was rotated by ~8° around its long axis with its shoulder side moving closer to the large subunit while its platform side moving away from it. **(B)** Superimposition of the Class I cryo-EM map with the factor-free 55S mitoribosome (light brown) (Amunts et al., 2015) revealed an overall ~8.5° rotation of the 28S subunit (yellow) in an anti-clockwise direction relative to the 39S subunit (blue). Additionally, the head domain of the 28S subunit rotated by ~4° towards the tRNA exit (E) site in a roughly orthogonal direction to the inter-subunit motion. In both the panels, landmarks of the 28S subunit: h, head; b, body. Landmarks of the 39S subunit: CP, central protuberance; Sb, stalk base; L1, MRP uL1m.

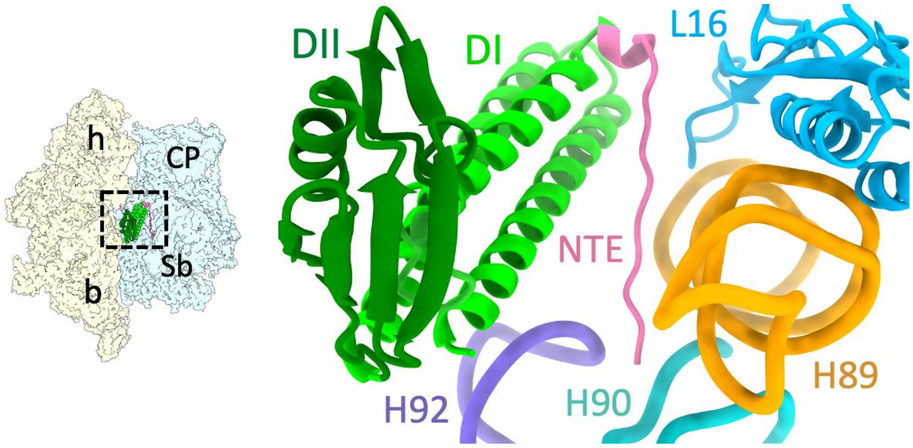


**Fig. S4. Interactions of $RRF_{mt}$ NTE with the mitoribosomal components. (A)** Simultaneous interactions of $RRF_{mt}$ NTE (pink) of $RRF_{mt}$ with various functionally important helices of the 16S rRNA such as H89 (orange), H90 (turquoise), H92 (purple) and MRP uL16m (light blue). Thumbnail to the left depicts an overall orientation of the 55S mitoribosome, with semitransparent 28S (yellow) and 39S (blue) subunits, and overlaid positions of ligands. Landmarks on the thumbnail: h, head, and b, body of the 28S subunit, and CP, central protuberance; Sb, stalk base of the 39S subunit.

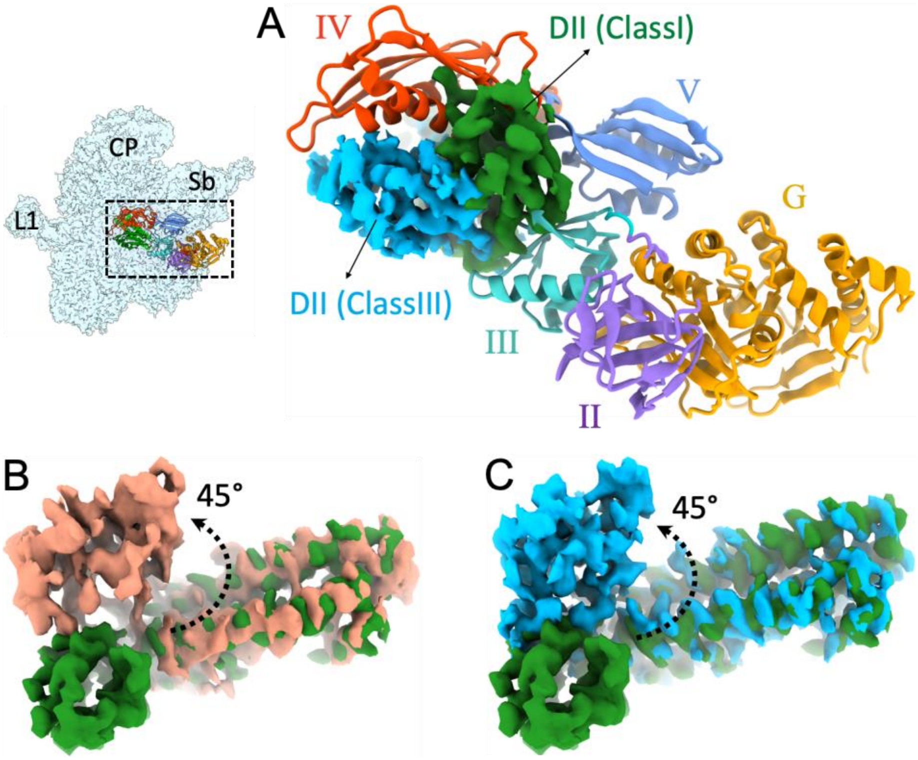


**Fig. S5. Domain II of RRF$_{mt}$ undergoes large conformational change to avoid steric clash with EF-G2$_{mt}$. (A)** Superposition of the 55S•RRF$_{mt}$ complex (Class I) with the 39S•RRF$_{mt}$•EF-G2$_{mt}$•GMPPCP complex (Class III) shows that the orientation of domain II (green) in Class I complex would prevent the binding of EF-G2$_{mt}$ to the RRF$_{mt}$-bound 55S mitoribosome due to direct steric conflict with the EF-G2$_{mt}$'s domains III (cyan), IV (red) and V (blue). **(B, C)** comparison of orientations of RRF$_{mt}$ domain II in Class I (green) Class II (light brown) and Class III (light blue) complexes. Thumbnail to the top left depicts an overall orientation of the 39S subunit (semitransparent blue) with overlaid positions of the ligands for panel A. Landmarks on the thumbnail: CP, central protuberance; Sb, stalk base; L1, MRP uL1m.

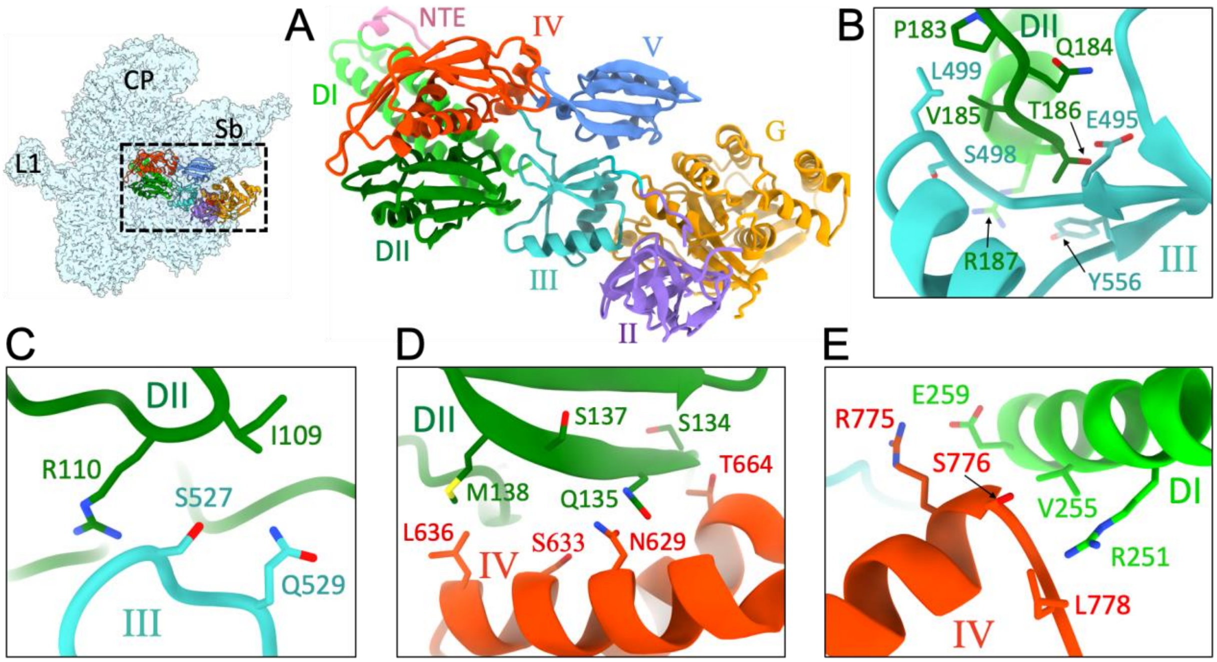


**Fig. S6. Interactions between RRF$_{mt}$ and EF-G2$_{mt}$ in the Class III complex. (A)** Domain II of RRF$_{mt}$ (dark green) is positioned in the pocket formed between domains III (cyan), IV (red) and V (blue) of EF-G2$_{mt}$. Thumbnail to the left of panel A depicts an overall orientation of the 39S subunit (semitransparent blue) with overlaid positions of the ligands. Landmarks on the thumbnail: CP, central protuberance; Sb, stalk base; L1, MRP uL1m. **(B, C)** Interactions of RRF$_{mt}$ domain II with domain III of EF-G2$_{mt}$. **(D)** Interactions of RRF$_{mt}$ domain II with domain IV of EF-G2$_{mt}$. **(E)** Interactions of RRF$_{mt}$ domain I with domain IV of EF-G2$_{mt}$.

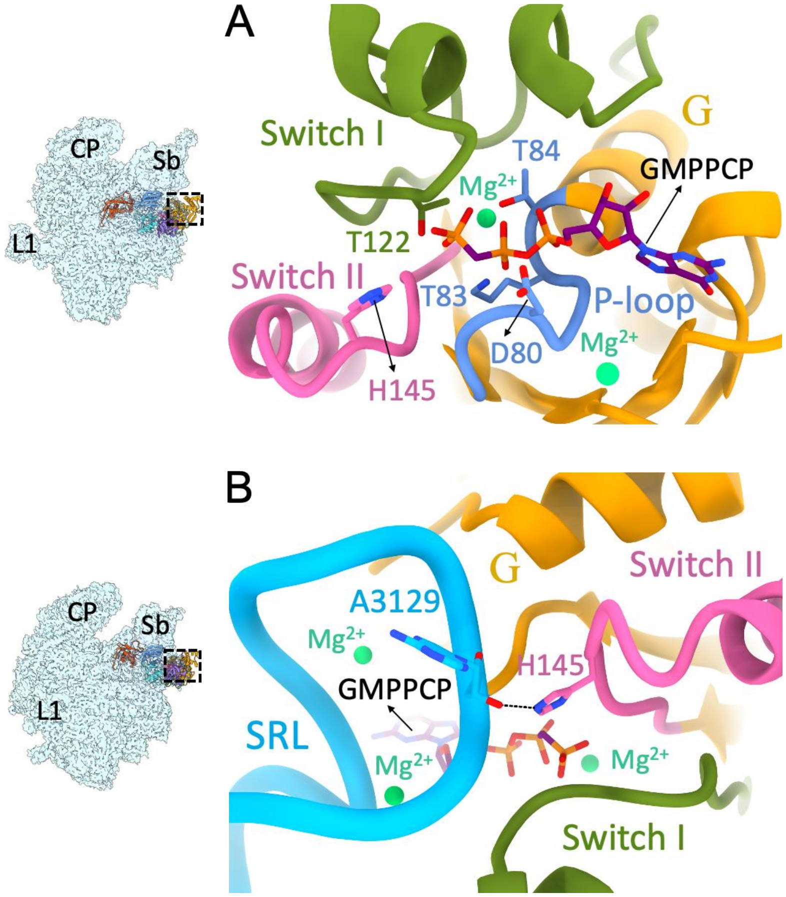


**Fig. S7. Interactions of the functionally essential elements within the G domain of EF-G2$_{mt}$ with GMPPCP and the SRL.** (**A**) GMPPCP is stably held in the nucleotide binding pocket through multiple interactions with conserved aa residues of the functionally important elements of the G domain such as Switch I (green), Switch II (pink) and the P-loop (blue). Magnesium ions present in the vicinity are shown as light green spheres. **(B)** The conserved H145 residue that is known to play a central role during GTP hydrolysis (Tourigny et al., 2013) is stabilized in its active conformation by interacting with the sugar moiety of the highly conserved A3129 residue from the SRL. Colors of the G domain components in the panels **A** and **B** are matched. Thumbnails to the left depict overall orientations of the 39S subunit (semitransparent blue) and overlaid ligands. Landmarks on the thumbnail: CP, central protuberance; Sb, stalk base; L1, MRP uL1m.

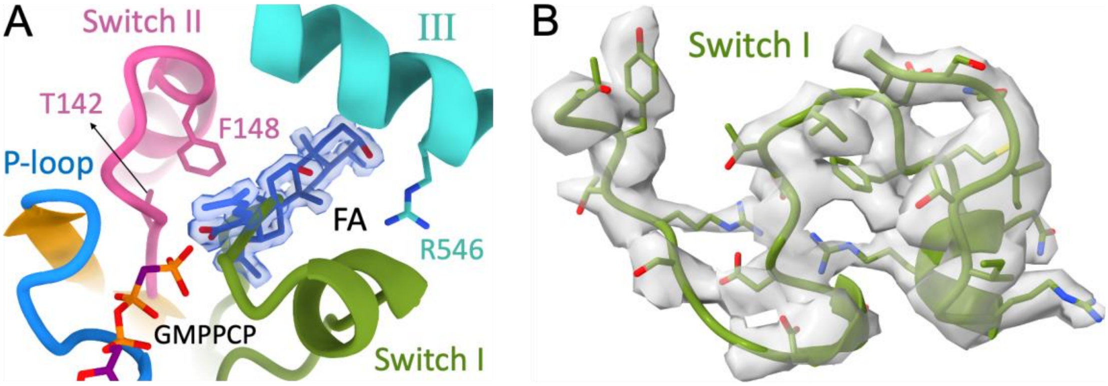


C

```
EF-G:       TAAVTTCFWKDHRINIIDTPGHVDFTIEVERSMRVLDGAIVV   107
EF-G1mt:    QSAATYTMWKDVNINIIDTPGHVDFTIEVERALRVLDGAVLV   144
EF-G2mt:    QSAAVTFDWKGYRVNLIDTPGHVDFTLEVERCLRVLDGAVAV   165

EF-G:       THPETGQTIISGMGELHLEIIVDRLKREFKVDANVGKPQVAY   483
EF-G1mt:    FDTENKETVISGMGELHLEIYAQRLEREYGCPCITGKPKVAF   525
EF-G2mt:    LDPDSGQTVLCGMGELHIEIIHDRIKREYGLETYLGPLQVAY   564
```

**Fig. S8. Stabilized Switch I region in EF-G2$_{mt}$ prevents FA from accessing its binding site.** **(A)** Superimposition of the FA molecule (dark blue) from the bacterial 70S•EF-G•GDP•FA complex (Gao et al., 2009) into the G domain of EF-G2$_{mt}$ in our 39S•RRF$_{mt}$•EF-G2$_{mt}$•GMPPCP complex reveals that the putative FA binding site is sterically blocked by the switch I region. Key aa residues known to be important for the stable binding of FA are shown with sidechains. Colors of the G domain components are similar as in Fig. S7. **(B)** Cryo-EM density corresponding to the switch I region (green) extracted from the 39S•RRF$_{mt}$•EF-G2$_{mt}$•GMPPCP complex. **(C)** The aa residues that are known to be necessary for the stable binding of FA (Gao et al., 2009) are conserved in all the three EF-Gs and are highlighted in pink.

**Table S1.** Data collection, Refinement and Model validation.

| Description | 55S•$RRF_{mt}$ (Class I) | 55S•$RRF_{mt}$•EF-$G2_{mt}$ (Class III) |
|---|---|---|
| **Data collection and Refinement** | | |
| Microscope | FEI Titan Krios | |
| Voltage (kV) | 300 | |
| Pixel size (Å) | 1.073 | |
| Defocus range (μm) | 1.0 to 3.0 | |
| Average $e^-$ dose per image ($e^-$/Å$^2$) | 71.6 | |
| Software | cryoSPARC | |
| Particles (initial) | 1,140,751 | |
| Particles (final) | 93,212 | 132,008 |
| Symmetry | C1 | C1 |
| FSC-threshold | 0.143 | 0.143 |
| Resolution (Å) | 3.49 | 3.15 |
| Map-sharpening *B* factor (Å$^2$) overall | 51.4 | 55.6 |
| **RMS deviations** | | |
| Bonds (Å) | 0.00 | 0.01 |
| Angles (°) | 0.05 | 0.05 |
| **Molprobity clashscore** | 1.97 (77$^{nd}$) | 2.03 (74$^{th}$) |
| Clashscore, all atoms | 10.33 | 10.97 |
| **Rotamer outliers (%)** | 0.68 | 0.84 |
| **Ramachandran plot** | | |
| Favored (%) | 93.24 | 92.23 |
| Outliers (%) | 0.19 | 0.50 |
| **RNA** | | |
| Correct sugar puckers (%) | 98.42 | 98.17 |
| Angle outliers (%) | 0.01 | 0.01 |
| Bond outliers (%) | 0.00 | 0.00 |
| Good backbone conformations (%) | 76.24 | 77.12 |
| **Model composition** | | |
| RNA bases<br>Protein residues | 2,527<br>14,369 | 1,583<br>9,322 |
| **Accession codes** | | |
| Cryo-EM maps | EMD-23096 | EMD-23121 |
| PDB ID | 7L08 | 7L20 |

### Supplemental References


Adams, P.D., Afonine, P.V., Bunkoczi, G., Chen, V.B., Davis, I.W., Echols, N., Headd, J.J., Hung, L.W., Kapral, G.J., Grosse-Kunstleve, R.W., *et al.* (2010). PHENIX: a comprehensive Python-based system for macromolecular structure solution. Acta Crystallogr D Biol Crystallogr *66*, 213-221.

Amunts, A., Brown, A., Toots, J., Scheres, S.H.W., and Ramakrishnan, V. (2015). Ribosome. The structure of the human mitochondrial ribosome. Science *348*, 95-98.

Chen, V.B., Arendall, W.B., 3rd, Headd, J.J., Keedy, D.A., Immormino, R.M., Kapral, G.J., Murray, L.W., Richardson, J.S., and Richardson, D.C. (2010). MolProbity: all-atom structure validation for macromolecular crystallography. Acta Crystallogr D Biol Crystallogr *66*, 12-21.

Emsley, P., Lohkamp, B., Scott, W.G., and Cowtan, K. (2010). Features and development of Coot. Acta Crystallogr D Biol Crystallogr *66*, 486-501.

Gao, Y.G., Selmer, M., Dunham, C.M., Weixlbaumer, A., Kelley, A.C., and Ramakrishnan, V. (2009). The structure of the ribosome with elongation factor G trapped in the posttranslocational state. Science *326*, 694-699.

Koripella, R.K., Sharma, M.R., Bhargava, K., Datta, P.P., Kaushal, P.S., Keshavan, P., Spremulli, L.L., Banavali, N.K., and Agrawal, R.K. (2020). Structures of the human mitochondrial ribosome bound to EF-G1 reveal distinct features of mitochondrial translation elongation. Nat Commun *11*, 3830.

Koripella, R.K., Sharma, M.R., Risteff, P., Keshavan, P., and Agrawal, R.K. (2019). Structural insights into unique features of the human mitochondrial ribosome recycling. Proc Natl Acad Sci U S A *116*, 8283-8288.

Mondal, S., Hsiao, K., and Goueli, S.A. (2015). A Homogenous Bioluminescent System for Measuring GTPase, GTPase Activating Protein, and Guanine Nucleotide Exchange Factor Activities. Assay Drug Dev Technol *13*, 444-455.

Pettersen, E.F., Goddard, T.D., Huang, C.C., Couch, G.S., Greenblatt, D.M., Meng, E.C., and Ferrin, T.E. (2004). UCSF Chimera--a visualization system for exploratory research and analysis. J Comput Chem *25*, 1605-1612.

Punjani, A., Rubinstein, J.L., Fleet, D.J., and Brubaker, M.A. (2017). cryoSPARC: algorithms for rapid unsupervised cryo-EM structure determination. Nat Methods *14*, 290-296.

Rohou, A., and Grigorieff, N. (2015). CTFFIND4: Fast and accurate defocus estimation from electron micrographs. J Struct Biol *192*, 216-221.

Roy, A., Kucukural, A., and Zhang, Y. (2010). I-TASSER: a unified platform for automated protein structure and function prediction. Nat Protoc *5*, 725-738.

Tourigny, D.S., Fernandez, I.S., Kelley, A.C., and Ramakrishnan, V. (2013). Elongation factor G bound to the ribosome in an intermediate state of translocation. Science *340*, 1235490.
